# A non-canonical EZH2/TRIM28 epigenetic axis drives heparan sulfate remodeling and melanoma metastasis

**DOI:** 10.64898/2026.01.21.700969

**Authors:** Neil G. Patel, Alexandra Drakaki, Farhan Valummel, Jack C. Moore, Amrita Basu, Bin Hu, Xiaolin Dong, Peng Zhao, Mees Botman, Emily Spector, Mark C. Mochel, Lance Wells, Jennifer E. Koblinski, Marten A. Hoeksema, Ryan J. Weiss

**Affiliations:** Complex Carbohydrate Research Center, University of Georgia, Athens, Georgia 30602, United States; Department of Biochemistry and Molecular Biology, University of Georgia, Athens, Georgia 30602, United States; Department of Medical Biochemistry, Amsterdam University Medical Center, University of Amsterdam, Amsterdam, The Netherlands; Massey Cancer Center, Virginia Commonwealth University; Richmond, VA, United States; Department of Pathology, Virginia Commonwealth University; Richmond, VA, United States

**Author notes:** To whom correspondence should be sent: Ryan J. Weiss, Ph.D., Complex Carbohydrate Research Center, University of Georgia, 315 Riverbend Road, Athens, GA 30602, Phone: 706/542-6445 FAX: 706/542-4412.

**Keywords:** glycosaminoglycans, heparan sulfate, epigenetic regulation, transcription, SULF1, EZH2, TRIM28, melanoma, metastasis

## Abstract

Melanoma progression and metastasis are driven not only by oncogenic alterations but also by epigenetic programs that dynamically remodel the tumor microenvironment. Heparan sulfate (HS) proteoglycans are key extracellular matrix components that integrate growth factor signaling, cell-matrix interactions, and migratory behavior by controlling ligand availability and receptor engagement, yet how chromatin-associated factors regulate HS remodeling in cancer remains poorly defined. Here, we identify the histone methyltransferase EZH2 as a key regulator of HS biosynthesis in melanoma. Integrated bioinformatic and genomic analyses revealed enrichment of EZH2 and additional Polycomb Repressive Complex (PRC) factors at regulatory regions of HS biosynthetic genes. CRISPR-mediated loss of EZH2 altered expression of multiple HS-modifying enzymes, most notably the secreted endosulfatases SULF1 and SULF2, resulting in enhanced HS 6-*O* sulfation and altered ligand binding at the cell surface. Unexpectedly, EZH2 promoted *SULF1* expression through a methyltransferase-independent mechanism via a non-canonical interaction with TRIM28, whereas *SULF2* was regulated through canonical PRC2-mediated repression. Functionally, SULF1 depletion impaired melanoma cell migration and invasion *in vitro* and reduced spontaneous metastasis in an orthotopic xenograft model. Together, these findings define an epigenetic axis linking chromatin regulation to extracellular glycan remodeling and identify HS-modifying enzymes as candidate targets to limit melanoma metastasis.

## INTRODUCTION

Melanoma is an aggressive form of skin cancer whose lethality stems primarily from its ability to rapidly metastasize. Although advances in targeted therapies and immunotherapies have improved outcomes, metastatic melanoma remains largely incurable^1^. When diagnosed at a localized stage, the five-year survival rate for melanoma exceeds ∼99%, but once the disease becomes metastatic, the five-year survival rate drops to ∼35%^1, 2^. Progression from a proliferative to an invasive state represents a defining transition in melanoma biology and is driven not only by genetic alterations but also by dynamic remodeling of the tumor microenvironment^3, 4^. In particular, remodeling of the extracellular matrix (ECM) plays a central role in enabling melanoma cell plasticity, invasion, and metastatic spread by shaping growth factor availability, receptor signaling, and cell-matrix interactions^5^.

Among the major molecular components of the ECM, heparan sulfate proteoglycans (HSPGs) are central mediators of cell-cell and cell-matrix communication and are increasingly recognized as key determinants of tumor behavior. HSPGs are core proteins covalently linked to long, linear heparan sulfate (HS) polysaccharides that consist of repeating disaccharides of glucosamine (GlcN) and uronic acids (GlcA, IdoA) that are heterogeneously sulfated. The sulfated domains dictate binding sites for a diverse set of protein ligands, including fibroblast growth factors (FGFs), vascular endothelial growth factors (VEGFs), WNTs, and chemokines (Fig. 1A)^6, 7^. Importantly, the fine structure of HS, defined by the spatial organization and density of sulfated domains, critically modulates downstream signaling and thereby influences processes such as proliferation, angiogenesis, and invasion^8^. HS sulfation patterning is established and dynamically regulated by a network of Golgi-localized 2-*O*, 3-*O*, and 6-*O* sulfotransferases and secreted extracellular sulfatases, including SULF1 and SULF2, which selectively remove 6-*O* sulfate groups from HS at the cell surface to modulate ligand-receptor interactions^9^. Dysregulation of HS biosynthetic and remodeling machinery is a recurring feature of cancer and has been directly implicated in melanoma progression^10, 11^. For example, distinct modes of HS remodeling have been implicated in melanoma progression, including HS chain cleavage by the endoglycosidase heparanase, which enhances melanoma cell migration and angiogenesis^12^, and alterations in HS sulfation modulate Wnt5A-driven invasive signaling programs characteristic of metastatic melanoma cells^13^. Genetic targeting of HS sulfation has also been shown to re-sensitize drug resistant melanoma cells to MAPK pathway inhibitors^14^. In addition, aberrant expression of specific HSPG core proteins, such as perlecan and syndecan- 1, correlates with metastatic lesions and melanoma invasiveness^15, 16^. Collectively, these findings establish HS structure and remodeling as critical regulators of the tumor microenvironment and melanoma progression. However, despite the clear functional importance of HS-modifying enzymes, the transcriptional and epigenetic mechanisms regulating their expression remain poorly understood.

**Figure 1.**
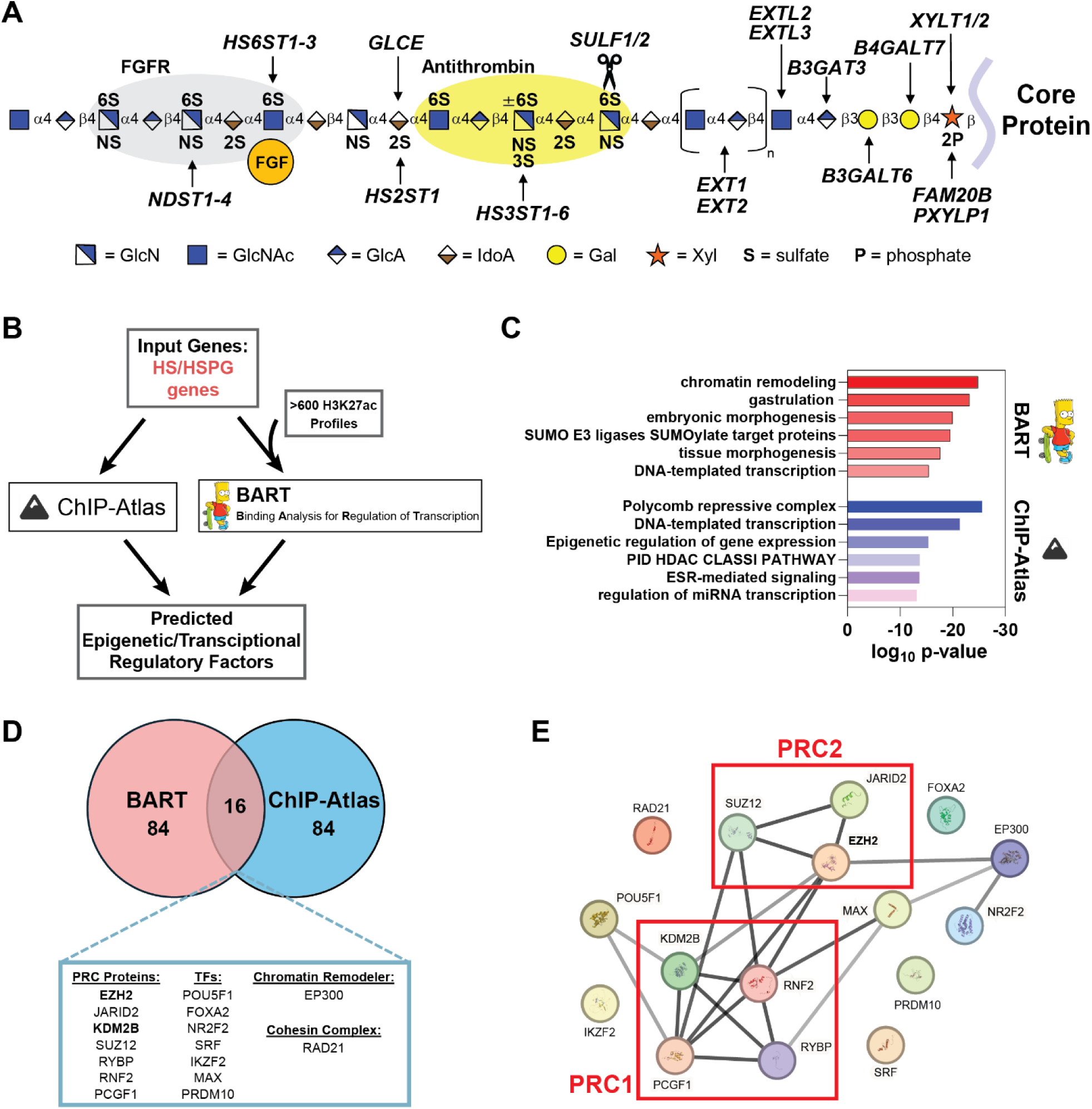
Dual bioinformatic screening approach uncovers predicted regulators of heparan sulfate biosynthesis. **(A)** Schematic representation of heparan sulfate (HS) biosynthesis. HS assembles while attached via a tetrasaccharide to a proteoglycan core protein. Secreted endosulfatases (SULF1/SULF2) are involved in the removal of 6-*O* sulfate groups. **(B)** Workflow for dual bioinformatic screening approach utilizing BART and ChIP-Atlas to mine publicly available genomic datasets for factors enriched at cis-regulatory regions of HS and HSPG genes. (**C**) Metascape pathway analysis of the top 100 unique hits from BART and ChIP-Atlas analyses, respectively. **(D)** Venn diagram of the intersection of the top 100 enriched factors from BART and ChIP-Atlas analyses. The 16 overlapping hits were characterized based on function. **(E)** STRING-DB analysis displaying the protein-protein interaction network of the overlapping identified factors reveals enrichment of Polycomb repressive complex (PRC) proteins.

Recent studies, including work from our group, have begun to reveal how HS biosynthesis, despite occurring in a non-template-driven manner^7^, is integrated into transcriptional and epigenetic gene regulatory networks. The zinc finger transcription factor ZNF263 was shown to repress genes encoding specific HS biosynthetic enzymes, acting as a central transcriptional regulator of heparin assembly and 3-*O* sulfation^17^. The histone demethylase KDM2B was identified as a chromatin-level modulator of HS biosynthesis and extracellular matrix genes, linking epigenetic control to structural and functional remodeling of the tumor microenvironment^18^. More recently, we demonstrated that the transcription factor TFCP2 represses expression of the HS endosulfatase, SULF1, leading to modulation of HS structure and melanoma cell growth^19^. Complementary studies have also highlighted the role of DNA methylation in cancer progression via transcriptional silencing of multiple HS biosynthetic enzymes, including exostosin-1 (EXT1)^20^, 3-*O* and 6-*O* sulfotransferases (HS3STs/HS6STs)^21, 22^, among others^23^. Together, these studies indicate that HS biosynthesis is tightly regulated by transcriptional and epigenetic mechanisms, yet the upstream oncogenic regulators that integrate these programs during melanoma progression remain largely undefined.

Given the critical importance of HS composition in modulating tumor cell signaling and extracellular interactions, identifying epigenetic and transcriptional regulators of HS-modifying enzymes would reveal new drug targets and key mechanisms driving melanoma invasion and metastasis. Here, we identify the histone methyltransferase Enhancer of Zeste Homolog 2 (EZH2) as a key upstream regulator of HS remodeling in melanoma. While EZH2 is a well-established oncogenic driver in melanoma^24, 25^, its role in regulating ECM composition and glycan presentation remains largely unexplored. We demonstrate that EZH2 alters HS composition and growth factor interactions at the cell surface via a methyltransferase-independent mechanism by controlling expression of the extracellular endosulfatase SULF1 through a non-canonical complex with the nuclear scaffolding protein, TRIM28. Disruption of this EZH2/TRIM28/SULF1 axis impairs cell migration and invasion, revealing a previously unrecognized epigenetic mechanism controlling HS remodeling during tumor progression. Moreover, depletion of SULF1 in human melanoma cells significantly reduced spontaneous metastases in an orthotopic xenograft model, directly implicating HS remodeling in melanoma metastatic behavior.

## RESULTS

### Bioinformatic analyses identify EZH2 as a regulator of HS biosynthesis

Heparan sulfate is a critical regulator of tumor proliferation, adhesion, and signaling, yet the molecular mechanisms controlling its biosynthesis and function are largely unknown. To identify upstream regulators of HSPG and HS biosynthesis, we analyzed cis-regulatory regions of all genes involved in HSPG assembly using two complementary bioinformatic tools, BART (Binding Analysis for Regulation of Transcription)^26^ and ChIP-Atlas^27^. Unlike traditional motif-based approaches, both BART and ChIP-Atlas directly utilize empirical publicly available ChIP-seq datasets to provide robust predictions of regulatory factors involved in gene expression changes. To identify potential regulatory factors, we utilized both tools in parallel to enhance confidence in uncovering relevant factors implicated in HS assembly (Fig. 1B). To benchmark this approach, we first input known genes involved in cholesterol biosynthesis, a well characterized metabolic pathway. BART and ChIP-Atlas analyses revealed multiple known regulators of cholesterol biosynthesis in the overlapping hits, including master regulators SREBF1/SREBF2, CREBBP, and EP300^28, 29^ (Supp. Fig. 1). Next, we analyzed an input gene list consisting of all genes encoding known HSPGs and HS biosynthetic enzymes^30^ using BART and ChIP-Atlas (Fig. 1A). Both tools gave results that converged on a shared set of regulatory factors enriched at HS gene loci, with strong enrichment of polycomb proteins, other chromatin modifiers, and transcription factors involved in cell differentiation and development (Fig. 1C). We prioritized the top 100 unique significantly enriched factors from both bioinformatic analyses, which revealed 16 shared predicted regulators, many of which are members of the Polycomb Repressive Complex 1 (PRC1) and PRC2, which are chromatin remodeling complexes with important roles in gene regulation^31^ and tumorigenesis^32^ (Fig. 1D). String-DB was utilized to visualize protein-protein interactions between the top 16 common hits, which confirmed that multiple identified factors physically interact and are in PRC1/PRC2 complexes (Fig. 1E). Notably, the histone demethylase and non-canonical PRC1 member, KDM2B, was included in the enriched overlapping hits, which we recently reported as a novel regulator of HS biosynthesis^18^, thus further validating our approach. Among the PRC-associated hits, the histone methyltransferase EZH2 emerged as the top enriched factor, ranked by significance across both datasets, and was prioritized due to its catalytic role in H3K27 methylation and established oncogenic functions in melanoma^24^. Somatic activating mutations and aberrant expression of EZH2 have implicated this epigenetic factor in melanoma progression and metastasis, where excessive H3K27me3 methylation on tumor suppressor genes can drive tumor progression and metastasis^25, 33^. It remains unclear how alterations in EZH2 expression and activity may impact HSPG structure and function in the tumor microenvironment and alter downstream cell signaling and tumorigenesis.

### Ablation of *EZH2* leads to alterations in HS structure and function

Given the established role of EZH2 in melanoma progression, we aimed to investigate its potential regulatory function in HS biosynthesis. To do this, we targeted *EZH2* via CRISPR/Cas9 in A375 human melanoma cells to generate a clonal *EZH2* knockout cell line (*EZH2^-/-^*). Successful gene knockout was confirmed by Western blot (Fig. 2A) and Sanger sequencing with no obvious morphological defects compared to wild-type (Suppl. Fig. 2A-B). To determine changes in HS composition and abundance, we isolated cell surface HS by trypsin digestion and anion exchange chromatography from wild-type and *EZH2^-/-^* cells. Purified HS was then depolymerized into disaccharides using heparin lyases, and the resulting disaccharides were subsequently analyzed by liquid chromatography and quantitative high-resolution mass spectrometry (LC-MS), using published methods^34, 35^. Interestingly, EZH2 loss resulted in an ∼2-fold increase in total cell surface HS abundance and redistribution of sulfation toward more highly sulfated domains, with a significant reduction in D0A6 (ΔUA-GlcNAc6S) and increased D2S6 (ΔUA2S-GlcNS6S) disaccharides (Fig. 2B-C, Suppl. Table 3). Quantification of sulfate groups per disaccharide confirmed a significant increase in tri-sulfated disaccharide species in *EZH2^-/-^* cells (Suppl. Fig. 2C, Suppl. Table 4). To examine potential alterations in HS chain length, wild-type and *EZH2^-/-^* cells were radiolabeled with ^35^SO_4_ for 24 hours at 37°C, and the [^35^S]HS was purified by anion exchange chromatography. Size exclusion chromatography of [^35^S]HS from wild-type and *EZH2^-/-^*cells revealed a modest reduction in HS chain length in *EZH2^-/-^* cells, with average estimated chain lengths of ∼35 and ∼30 kDa in wild-type and knockout cells, respectively (Fig. 2D).

**Figure 2.**
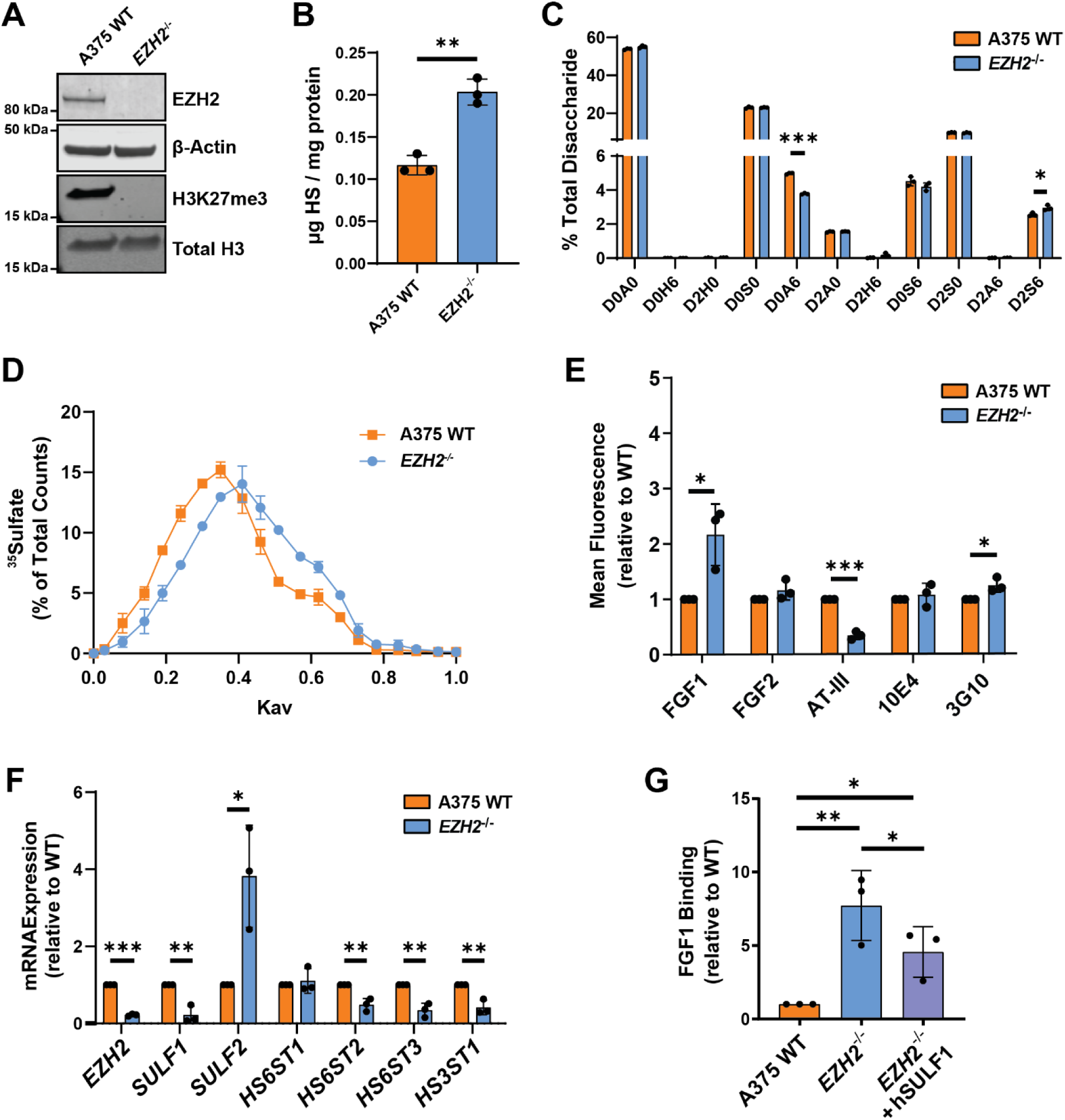
Targeting *EZH2* alters HS structure, abundance, and functional properties in human melanoma cells. **(A)** Western blot of EZH2 and H3K27me3 levels in A375 wild-type and *EZH2* knockout cells (*EZH2*^-/-^). **(B)** Liquid chromatography-mass spectrometry (LC-MS) quantification of total cell surface HS isolated from wild-type and *EZH2*^-/-^ cells. **(C)** LC-MS quantification of cell surface HS disaccharides in wild-type and *EZH2*^-/-^ cells. **(D)** Gel filtration chromatography (Sepharose CL-6B) of [^35^S]HS from wild-type and *EZH2*^-/-^ cells. **(E)** Cell surface binding assays of known HS-binding ligands, measured by flow cytometry. **(F)** qPCR mRNA expression analysis of a subset of genes encoding enzymes involved in 6-*O* and 3-*O* sulfation modification of HS. **(G)** Flow cytometry analysis of FGF1 binding for *EZH2*^-/-^ cells transfected with a human SULF1 plasmid relative to mock transfected control cells. Data are presented as mean ± SD (n = 3 independent experiments), ***p<0.001, **p<0.01, *p<0.05 by two-sided t-test.

To assess whether the observed compositional changes in HS derived from *EZH2^-/-^* cells could impact HS-protein interactions at the cell surface, we quantified cell surface binding of a panel of known HS-interacting proteins using flow cytometry. Notably, fibroblast growth factor 1 (FGF1) binding was significantly increased and anti-thrombin (AT-III) was drastically reduced in *EZH2^-/-^*cells (Fig. 2E). Interestingly, binding of FGF2 and an antibody recognizing *N*-sulfated hybrid domains of HS (10E4) remained unchanged (Fig. 2E), consistent with the minimal changes observed in 2-*O* and *N*-sulfated HS disaccharides in the LC-MS analysis (Fig. 2B). We also detected a slight increase in binding of an antibody that detects an HS neoepitope generated upon heparin lyase digestion (3G10) (Fig. 2E), which is consistent with an increase in cell surface HS-modified sites (Fig. 2B). The selective increase in FGF1 binding, together with unchanged FGF2 and 10E4 binding, is compatible with a redistribution of HS sulfation toward 6- *O*-sulfated domains^36, 37^. To corroborate these findings, we identified a second *EZH2^-/-^*knockout clonal line bearing different mutant alleles, which gave a similar binding profile (Suppl. Fig. 2D-F).

As EZH2 is a well-established chromatin remodeler and regulator of gene expression, we next investigated whether ablation of *EZH2* altered mRNA expression of key HS genes encoding HS modifying enzymes. We prioritized genes that were expressed in A375 wild-type cells^19^ and that are known to contribute to 6-*O*- and 3-*O*-sulfated residues important for FGF1 and AT-III binding^37, 38^, which were altered in *EZH2^-/-^*cells (Fig. 2). Due to the preferential recognition of 6-*O* sulfation by FGF1, we probed the expression of two extracellular HS 6-*O*-endosulfatases (*SULF1, SULF2*) and three HS 6-*O*-sulfotransferases (*HS6ST1, HS6ST2, HS6ST3*) by quantitative PCR. In addition, we probed the expression of the 3-*O*-sulfotransferase *HS3ST1*, which adds a rare 3-*O* sulfate group essential for AT binding^38^. Intriguingly, *SULF1* and *SULF2* were differentially altered upon *EZH2* inactivation, with a significant decrease in *SULF1* and concomitant increase in *SULF2* expression in *EZH2^-/-^* cells (Fig. 2F). *HS6ST2* and *HS6ST3* decreased in expression upon *EZH2* ablation. *HS3ST1* expression was also suppressed in *EZH2^-/-^* cells, consistent with the observed decrease in AT binding (Fig. 2E). Although expression of multiple HS sulfotransferase enzymes were altered in *EZH2^-/-^*cells, the dominant reduction in *SULF1* expression, previously shown to be the primary extracellular sulfatase expressed in A375 cells (Suppl. Fig. 2G)^18, 19^, suggested a central role for SULF1 in mediating the observed HS composition and ligand-binding phenotypes. To confirm this, we overexpressed human SULF1 in *EZH2^-/-^*cells, which led to a partial reversal of the enhanced FGF1 binding (Fig. 2G, Suppl. Fig. 2H), indicating that reduced SULF1 contributes significantly to the observed FGF1-HS binding increase upon ablation of EZH2.

### EZH2 regulates SULF1 and SULF2 via distinct regulatory mechanisms

Due to the established role of EZH2 as a canonical repressor within the PRC2 complex, we were intrigued by the opposing changes in *SULF1* and *SULF2* expression in *EZH2^-/-^* knockout cells. To investigate this further, we first aimed to determine if *SULF1* and *SULF2* are canonically regulated by the PRC2 complex by targeting two key PRC2 components, EED and SUZ12^32^, via siRNA-mediated knockdown. Upon confirmation of depletion of EED or SUZ12 in A375 wild-type cells (Fig. 3A, C), quantitative PCR analysis revealed *SULF1* expression was unchanged while *SULF2* expression was significantly elevated upon depletion of EED or SUZ12, respectively (Fig. 3B, D). Consistent with minimal effects on *SULF1* expression, PRC2 knockdown did not recapitulate the enhanced FGF1 binding observed in *EZH2*^-/-^ cells (Fig. 3E). These results suggested that *SULF2* is likely regulated by PRC2-mediated gene repression and does contribute to FGF1 binding, while the observed decrease in *SULF1* expression and increase in FGF1 binding in *EZH2^-/-^*cells was likely independent of canonical PRC2 activity.

**Figure 3.**
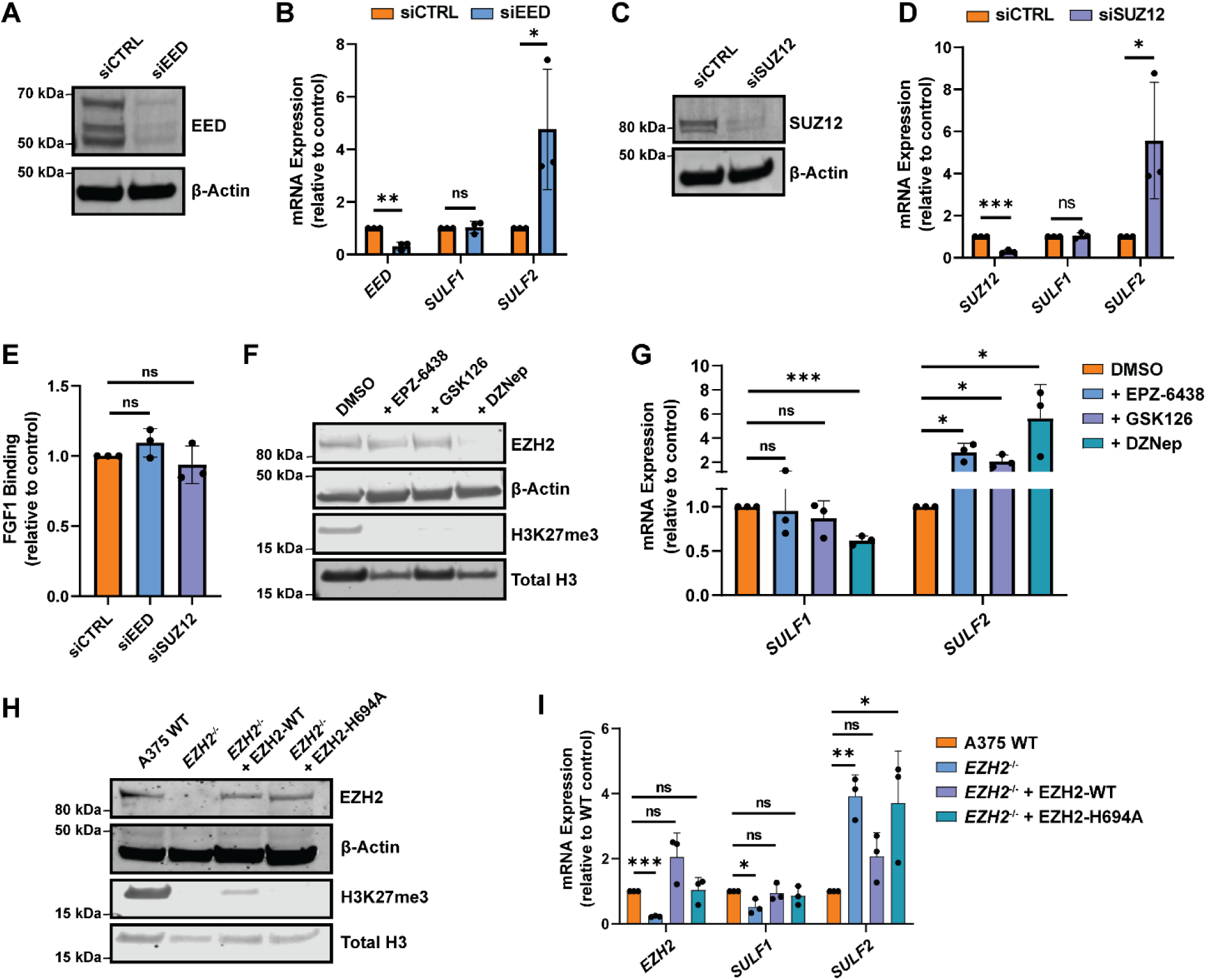
*SULF1* and *SULF2* are differentially regulated by EZH2 and the PRC2 complex in human melanoma cells. **(A)** Western blot validating transient siRNA-mediated knockdown of EED, a core PRC2 component, in A375 cells. **(B)** qPCR analysis of *SULF1* and *SULF2* mRNA expression upon knockdown of *EED* in A375 cells relative to scrambled siRNA control transfected cells. **(C)** Western blot validating transient siRNA-mediated knockdown of core PRC2 protein, SUZ12, in A375 cells. **(D)** qPCR analysis of *SULF1* and *SULF2* mRNA expression upon knockdown of *SUZ12* in A375 cells. **(E)** FGF1 cell surface binding assays upon knockdown of core PRC2 proteins, measured by flow cytometry. **(F)** Western blot analysis of EZH2 and H3K27me3 levels for A375 wild-type cells upon treatment with EZH2 inhibitors compared to DMSO (vehicle) treated control cells. EZH2 methyltransferase inhibitors (EPZ-6438 and GSK126) depleted global H3K27me3 levels without altering EZH2 protein levels, and DZNep promoted EZH2 protein degradation and depleted global H3K27me3 levels. **(G)** qPCR analysis probing *SULF1* and *SULF2* mRNA expression upon treatment with EPZ-6438, GSK126, or DZNep relative to vehicle-treated control cells. **(H)** Western blot confirming overexpression of EZH2 wild-type (EZH2-WT) and a catalytic mutant (EZH2-H694A) in *EZH2^-/-^* cells. **(I)** qPCR analysis of *SULF1* and *SULF2* expression upon rescue with EZH2-WT or EZH2-H694A relative to mock-transfected controls. Data are presented as mean ± SD (n = 3 independent experiments), ***p<0.001, **p<0.01, *p<0.05 by two-sided t-test.

To determine whether EZH2 catalytic activity mediates regulation of *SULF1* and *SULF2*, we treated A375 wild-type cells with diverse small molecule EZH2 inhibitors. EPZ-6438 and GSK126 were utilized to block EZH2 methyltransferase activity via competitive inhibition of the methyl donor S-adenosylmethionine (SAM)^39, 40^, whereas 3-Deazaneplanocin A (DZNep) treatment leads to depletion of EZH2 protein by inhibiting S-adenosyl-L-homocysteine hydrolase^41^. Treatment of A375 cells with EPZ-6438 or GSK126 (2 µM) for 72 hours resulted in depletion of global H3K27me3 levels without altering EZH2 protein abundance, as expected (Fig. 3F). Notably, catalytic inhibition led to an increase in *SULF2* expression but did not affect *SULF1* expression levels. In contrast, DZNep treatment decreased both EZH2 protein and H3K27me3 levels and resulted in reduced *SULF1* expression while also increasing *SULF2* (Fig. 3G). These findings indicate that maintenance of *SULF1* expression depends on EZH2 protein independent of its methyltransferase activity, whereas *SULF2* is repressed by EZH2 catalytic function, revealing mechanistically distinct regulatory modes for the two extracellular sulfatases.

To confirm these findings, we performed targeted EZH2 rescue experiments by transfecting *EZH2^-/-^* cells with CMV-driven vectors expressing either wild-type EZH2 (EZH2-WT) or a catalytic mutant (EZH2-H694A). Western blot analysis confirmed restoration of EZH2 protein levels in both EZH2-WT and EZH2-H694A transfected knockout cells, with partial rescue of H3K27me3 observed only in the EZH2-WT transfected cells (Fig. 3H). Reintroduction of EZH2-WT or EZH2-H694A rescued *SULF1* expression, while only overexpression of EZH2-WT reduced *SULF2* expression to wild-type levels (Fig. 3I). Together, these results indicate that *SULF2* is regulated via EZH2’s canonical methyltransferase activity within the PRC2 complex, whereas *SULF1* expression depends on EZH2 through a PRC2- and methylation-independent mechanism.

### *SULF1* is regulated by a non-canonical EZH2/TRIM28 axis

To investigate non-canonical mechanisms by which EZH2 regulates *SULF1*, we first mapped the EZH2 protein interactome in A375 wild-type cells. Endogenous EZH2 was immunoprecipitated and analyzed by liquid chromatography-tandem mass spectrometry using published methods, with minor modifications^42^. This analysis identified 141 interacting proteins, including multiple canonical PRC2 components (EZH2, EED, SUZ12, JARID2, RBBP4, RBBP7)^43^, validating the robustness of the approach. Among the additional interactors, we focused on four factors previously reported to associate with EZH2, including TRIM28^44^, MTF2^45^, DDX5^46^, and CTNNB1^47^, as identified by STRING database analysis (Fig. 4A). Notably, previous studies in breast cancer cells suggested EZH2’s interaction with TRIM28/KAP1, a multifunctional epigenetic scaffold protein, can positively regulate *SULF1* transcripts^44^, prompting us to examine whether this complex operates in melanoma cells. We validated that EZH2 and TRIM28 interact in A375 cells via immunoprecipitation of endogenous TRIM28 and blotting for EZH2, confirming that this interaction occurs under physiological expression conditions (Fig. 4B). Functionally, stable knockdown of *TRIM28* in A375 cells (shTRIM28) reduced *SULF1* mRNA levels to a degree comparable to EZH2 loss (Fig. 4C-D), phenocopying *EZH2^-/-^* cells and supporting a shared regulatory pathway controlling *SULF1* expression.

**Figure 4.**
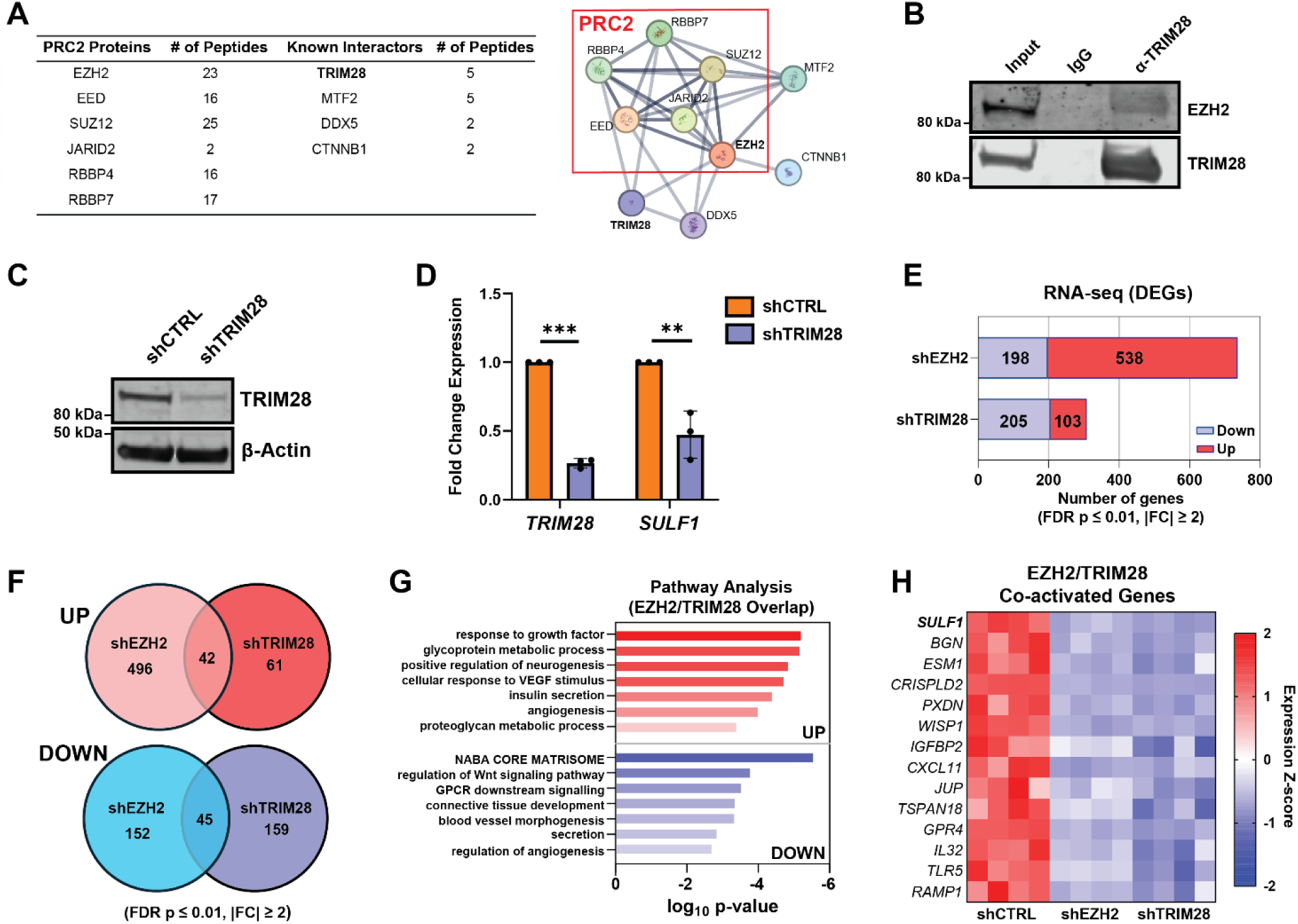
EZH2 and TRIM28 co-regulate *SULF1* and genes associated with the extracellular matrix and growth factor signaling. **(A)** Immunoprecipitation of EZH2 in A375 wild-type cells and LC-MS/MS proteomics reveals canonical PRC2 proteins and other known EZH2 interactors (table inset). STRING-DB analysis of top hits reveals the EZH2 interactome (right). **(B)** Endogenous TRIM28 in A375 cells was immunoprecipitated and immunoblotted to detect EZH2. **(C)** Western blot validating shRNA-mediated knockdown of *TRIM28* in A375 cells (shTRIM28). **(D)** qPCR analysis revealing that *SULF1* mRNA expression is decreased upon stable knockdown of *TRIM28*. **(E)** Number of DEGs in the RNA-seq data (FDR p-value ≤ 0.01, |FC| ≥ 2). The number of significantly up- and down-regulated genes for shEZH2 and shTRIM28 is included in the inset table. **(F)** Venn diagram showing overlap of up- and down-regulated genes in shEZH2 and shTRIM28 RNA-seq datasets (FDR p-value ≤ 0.01, |FC| ≥ 2). **(G)** Pathway enrichment analysis of overlapping DEGs between shEZH2 and shTRIM28 populations (FDR p-value ≤ 0.01, |FC| ≥ 2). **(H)** Heatmap of expression z-scores for a subset of co-activated DEGs shared between shEZH2 and shTRIM28 cells relative to shCTRL controls. Red represents higher expression and blue indicates lower expression. Data are presented as mean ± SD (n = 3 independent biological replicates, n = 4 independent biological replicates for RNA-seq experiments), ***p<0.001, **p<0.01, *p<0.05 by two-sided t-test.

To define the broader transcriptional programs co-regulated by EZH2 and TRIM28, we utilized shTRIM28 cells and generated an additional knockdown line targeting *EZH2* (shEZH2) (Suppl. Fig. 3A-B) and performed RNA sequencing. Differential expression analysis was conducted using a stringent threshold (FDR p-value ≤ 0.01; |fold change| ≥ 2). This analysis revealed that depletion of *EZH2* or *TRIM28* altered the expression of 737 and 308 genes, respectively, compared to a scrambled shRNA control cell line (shCTRL), indicating broad transcriptional remodeling. The majority (> 70%) of differentially expressed genes shEZH2 cells were upregulated, consistent with canonical PRC2-mediated gene repression, while shTRIM28 cells showed a larger subset (∼63%) of downregulated transcripts (Fig 4E). Pathway analysis of the significant DEGs overlapping across both cell lines revealed shared pathways related to cell adhesion, extracellular matrix, and processes involved in cell growth and activation (Suppl. Fig. 3C-D). Notably, we detected a significant overlap of 42 upregulated and 45 down-regulated DEGs that were shared between shEZH2 and shTRIM28 lines (Fig. 4F). These gene sets included factors involved in extracellular matrix remodeling, growth factor signaling, and proteoglycan-associated genes (Fig. 4H). Among the co-activated genes, *SULF1* emerged as one of the most significantly downregulated transcripts common to both knockdown conditions (Fig. 4G). Together, these results suggest a shared role for EZH2 and TRIM28 in coordinating broader ECM remodeling and signaling programs in melanoma.

To delineate the genomic mechanisms underlying differential regulation of *SULF1* and *SULF2*, we performed chromatin immunoprecipitation sequencing (ChIP-seq) to map the genomic localization of relevant interactors of EZH2 for canonical (EED, H3K27me3) versus non-canonical (TRIM28) activity. Globally, we found enrichment of TRIM28 and EED primarily at promoter regions near the TSS, while H3K27me3 marks were enriched at both promoters and within gene bodies, consistent with established PRC2 chromatin distributions (Fig. 5A-C). Specifically, we found that binding sites associated with canonical PRC2 activity (EED, H3K27me3) co-occupied the *SULF2* promoter and gene body, consistent with classical PRC2-mediated repression (Fig. 5D). In contrast, TRIM28 was enriched at the upstream regulatory region of *SULF1*, which lacked detectable EED or H3K27me3 signal (Fig. 5E). Globally, TRIM28 binding occurred at ∼9,090 sites at or near (within 10 kb) promoter regions and coding gene exons (Fig. 5F). To identify potential co-regulated target genes of TRIM28 in A375 cells, we integrated TRIM28 ChIP-seq data with our shTRIM28/shEZH2 RNA-seq data and found 9 genes, including *SULF1*, that were bound by TRIM28 and dependent on both TRIM28 and EZH2 for transcriptional activation (Fig. 5G). To confirm whether EZH2 localization at *SULF1* depends on TRIM28, we performed ChIP-qPCR for EZH2 in A375 wild-type and shTRIM28 cells. Quantification using primers targeting the *SULF1* upstream regulatory region revealed significantly reduced EZH2 occupancy in TRIM28-depleted cells (Fig. 5H), indicating that TRIM28 is critical for efficient EZH2 occupancy at the *SULF1* locus. Taken together, these findings support a model in which *SULF2* is regulated by canonical, methyltransferase-dependent PRC2 activity, whereas *SULF1* is controlled by a non-canonical EZH2/TRIM28 complex that directs EZH2 to the *SULF1* cis-regulatory region.

**Figure 5.**
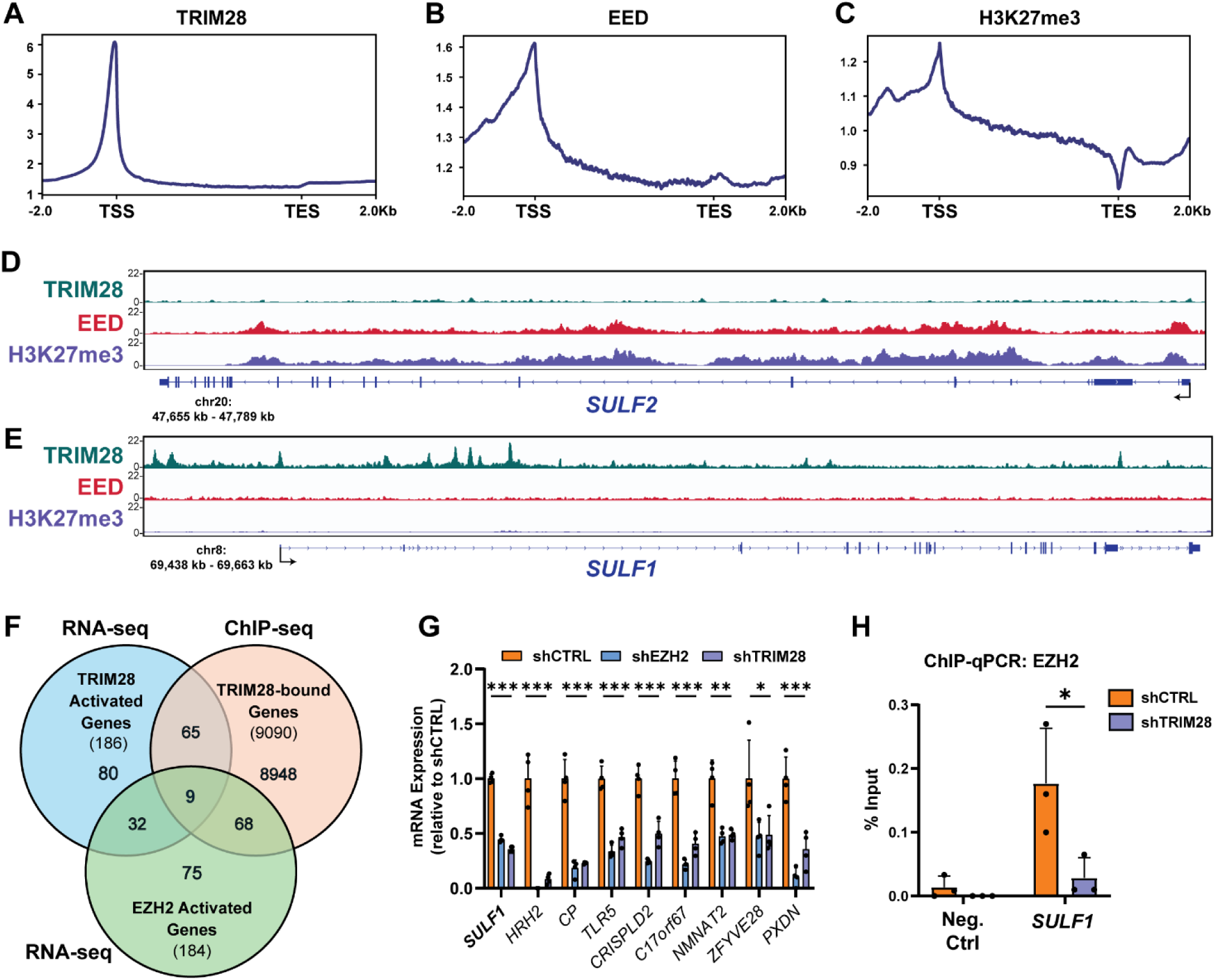
Distinct chromatin association patterns link EZH2/TRIM28 activity to *SULF1* activation and PRC2-mediated repression of *SULF2.* Profile plots showing genome-wide localization of **(A)** TRIM28, **(B)** EED, and **(C)** H3K27me3 relative to annotated gene bodies in A375 wild-type cells (hg38 genome, TSS: transcription start site, TES: transcription end site). Genome browser tracks showing TRIM28, EED, and H3K27me3 ChIP-seq signal at the *SULF2* (D) and *SULF1* (E) loci. **(F)** Venn diagram showing overlap between genes positively regulated by EZH2 and TRIM28 (RNA-seq) and genes bound by TRIM28 (ChIP-seq). **(G)** ChIP–qPCR assessing TRIM28-dependent enrichment of EZH2 at the *SULF1* locus in control and TRIM28-depleted A375 melanoma cells. Data are presented as mean ± SD (n = 3 independent biological replicates), ***p<0.001, **p<0.01, *p<0.05 by two-sided t-test.

### Targeting the EZH2/TRIM28/SULF1 axis impairs melanoma cell migration and invasion *in vitro*

RNA-seq analysis of shEZH2 and shTRIM28 knockdown lines revealed shared alterations in genes involved in cell adhesion and extracellular matrix organization (Suppl. Fig. 3D), suggesting that EZH2 and TRIM28 cooperatively regulate transcriptional programs associated with melanoma cell motility. Given that SULF1 was identified as a common downstream target of both factors and has been implicated in modulating tumor-associated signaling in other cancer contexts, we generated a stable *SULF1* knockdown cell line (shSULF1) to examine its contribution as an extracellular effector within this regulatory axis. shSULF1 knockdown cells exhibited reduced SULF1 expression and enhanced FGF1 binding, as expected (Suppl. Fig. 4A-B). Intriguingly, depletion of *EZH2, TRIM28,* or *SULF1* resulted in increased cell growth over time versus wild-type cells (Fig. 6A-B), yet all three knockdown lines exhibited a significant reduction in transwell migration (Fig. 6C-D) and invasion through Matrigel (Fig. 6E-F). Importantly, these findings were reproduced in an independent BRAF-mutant melanoma line, SKMEL-5, in which shRNA-mediated knockdown of *EZH2* or *SULF1* resulted in reduced migration and invasion compared to wild-type control cells (Suppl. Fig. 4C-J). Collectively, these data demonstrate that the EZH2/TRIM28/SULF1 axis promotes a migratory and invasive melanoma cell phenotype *in vitro*, while restraining proliferative capacity. This uncoupling of proliferation and invasion is consistent with established models of melanoma phenotypic plasticity^3^ and suggests that non-canonical EZH2-mediated regulation of heparan sulfate remodeling plays a critical role in controlling metastatic behaviors.

**Figure 6.**
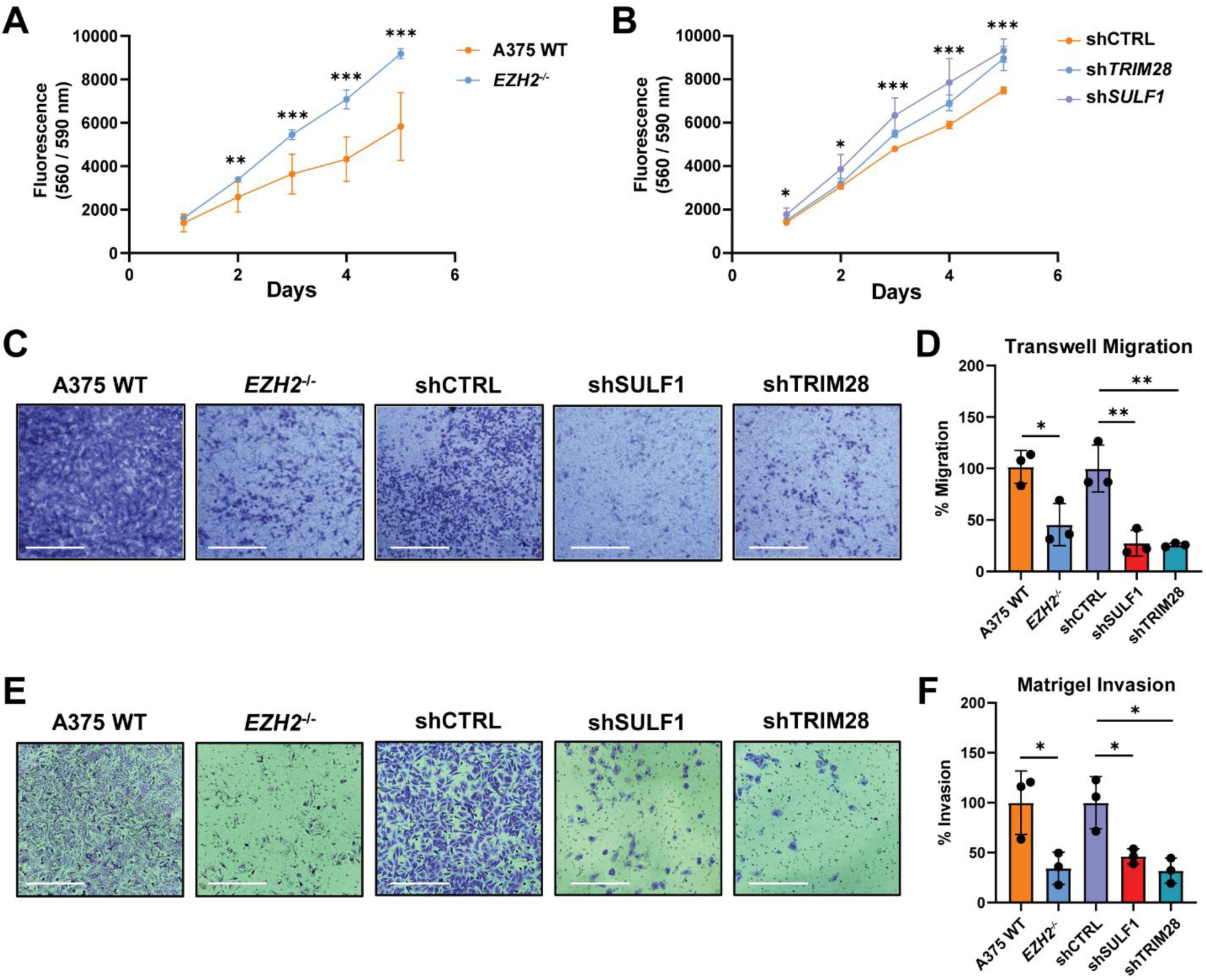
Targeting the EZH2/TRIM28/SULF1 axis reduces migration and invasion *in vitro.* **(A)** Cell growth assay (0-5 days) in A375 wild-type and *EZH2*^-/-^ cells. **(B)** Cell growth assay (0-5 days) in A375 shRNA-mediated knockdown cell lines (shCTRL, shSULF1, shTRIM28) relative to shCTRL cells. **(C)** Representative images of migrated cells stained with crystal violet (48-hour incubation). Scale bar: 360 µm. **(D)** Quantification of cell migration assays by measuring absorbance (590 nm) of crystal violet eluted from transwell inserts. Data was normalized to respective wild-type or shCTRL cells. **(E)** Representative images of Matrigel invading cells stained with crystal violet (36-hour incubation). Scale bar: 360 µm. **(F)** Quantification of cell invasion assays by measuring absorbance (590 nm) of crystal violet eluted from Matrigel inserts. Data was normalized to respective wild-type or shCTRL cells. All data are presented as mean ± SD (n = 3 independent biological replicates), ***p<0.001, **p<0.01, *p<0.05 by two-sided t-test.

### Depletion of SULF1 reduces melanoma metastasis *in vivo*

To assess the role of SULF1 in melanoma tumor growth and metastatic dissemination *in vivo*, we established a spontaneous metastasis xenograft model in NSG mice. First, A375 shCTRL and shSULF1 cells were engineered to stably express luciferase (shCTRL-Luc, shSULF1-Luc). Both cell lines exhibited comparable luminescence activity upon treatment with luciferin (Supp. Fig 5A). Cells were then injected orthotopically into the dermis of the right and left flanks of NSG mice to promote tumor formation and allow spontaneous metastases (Fig. 7A). Bilateral injections were performed to increase overall tumor burden per animal, given the modest metastatic potential of A375 cells^48^. SULF1 depletion resulted in a modest but statistically detectable difference in primary tumor growth trajectories over time, while the overall extent of primary tumor growth remained broadly comparable between groups (Fig. 7B). At the study endpoint, when total tumor burden approached ∼2 cm^3^ per mouse, IVIS imaging of resected tissues revealed spontaneous metastases in the lungs, liver, bone, and brain. Notably, mice bearing shSULF1 tumors exhibited a significant reduction in the frequency of distal metastases per mouse, with the most pronounced effect observed in the lungs and similar trends across other organs (Fig. 7C-D, Suppl. Fig. 5C-D). Chi-squared analysis confirmed a significant decrease in metastatic frequency across the cohort, with several mice harboring shSULF1 tumors displaying no detectable metastases. Histological analysis of primary tumors by H&E staining revealed expanded necrotic regions in shSULF1 tumors compared with shCTRL controls (Fig. 7E-F, Suppl. Fig. 5E), which could reflect altered tumor-stromal interactions and/or impaired vascular support. Together, these findings indicate that SULF1 plays a prominent role in promoting melanoma metastatic dissemination with relatively modest effects on primary tumor growth.

**Figure 7.**
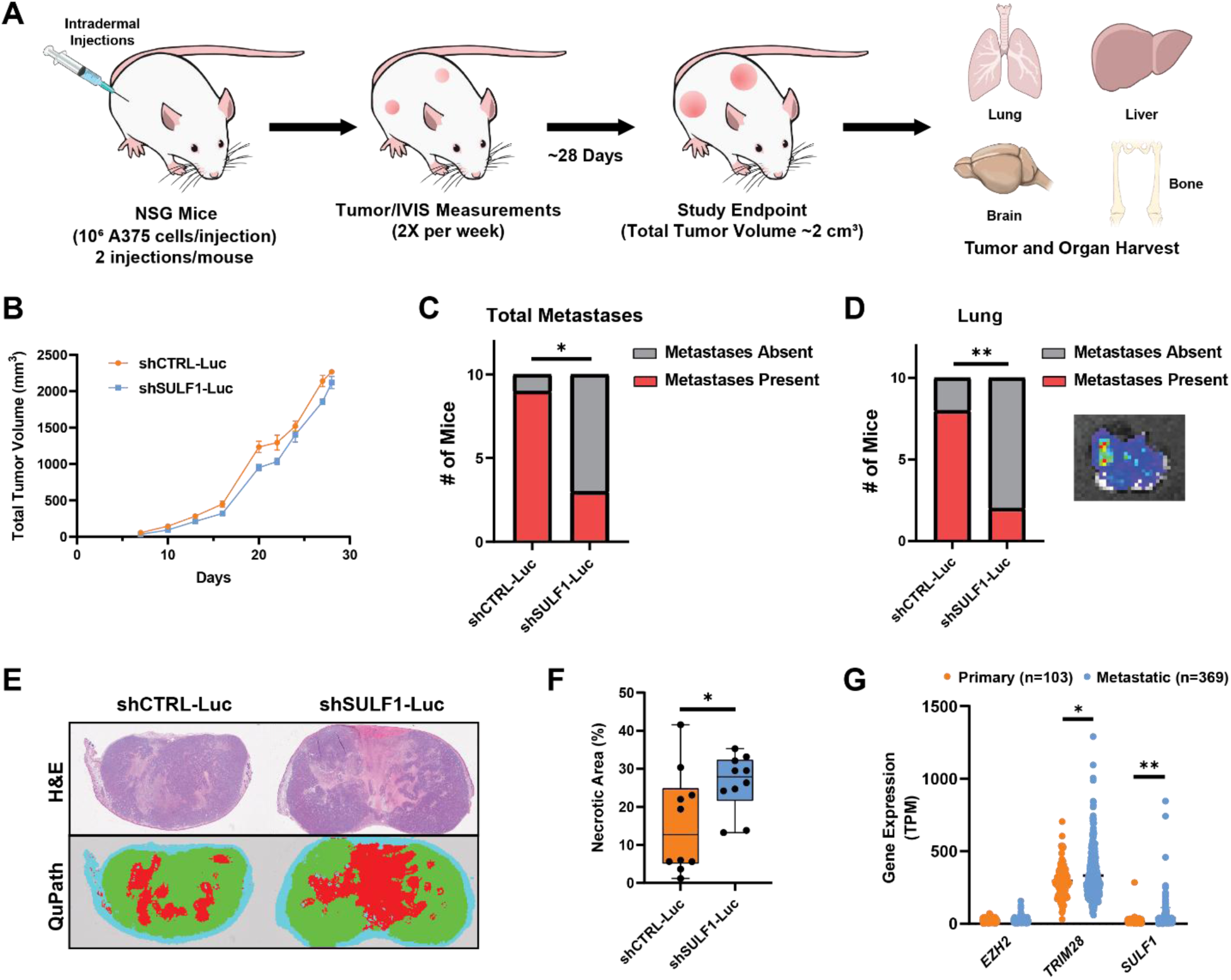
SULF1-deficient cells exhibit reduced metastasis in an orthotopic xenograft model. **(A)** Study design of an A375 melanoma xenograft model in NSG mice to assess primary tumor growth and metastatic dissemination of SULF1-deficient cells. **(B)** Total primary tumor volume measured longitudinally (mm³). Mice were euthanized upon reaching a humane endpoint defined as total tumor burden ≥ 2000 mm³, followed by excision of primary tumors and collection of organs for metastatic analysis (n = 10 mice per group; data are presented as mean ± SEM and represent total tumor volume per mouse, p = 0.0022). **(C)** Contingency analysis showing the number of mice with detectable metastases, as assessed by ex vivo bioluminescence imaging, in any of the indicated organs (lungs, liver, brain, bone, or lymph nodes) (n = 10 mice per group). **(D)** Contingency plot depicting the number of mice with observed metastases in the lung. Inset: representative image of a shCTRL-injected mouse lung showing luminescence signal from A375 cells is included as an inset. **(E-F)** Quantification of tumor necrosis expressed as necrotic area (red) relative to total tumor area, determined by H&E staining and QuPath and confirmed by a pathologist (n = 10 mice per group, data represent an average of total tumor necrotic area per mouse). **(G)** RNA-seq data from the TCGA-SKCM cohort comparing *EZH2*, *TRIM28*, and *SULF1* expression in primary melanoma tumors (n = 103) versus metastatic melanoma tumors (n = 369). Data are presented as mean ± SEM, ***p<0.001, **p<0.01, *p<0.05, unless otherwise stated above. Statistical significance was determined using two-sided tests as indicated (tumor growth: mixed-effects model for repeated measures; metastasis incidence: chi-squared test; tumor necrosis: two-sided t-test).

Finally, to evaluate the clinical relevance of the EZH2/TRIM28/SULF1 axis in human melanoma, we analyzed RNA-seq data from the TCGA skin cutaneous melanoma (SKCM) cohort. Expression of *TRIM28* and *SULF1* were significantly elevated in metastatic versus primary melanoma samples, whereas *EZH2* expression was not significantly altered (Fig. 7G). These clinical data are consistent with a role for TRIM28 and SULF1 in metastatic melanoma and support the functional relevance of this regulatory axis in human disease.

## DISCUSSION

In this study, we present an unexpected regulatory mechanism by which melanoma cells remodel their extracellular glycan landscape to promote invasive and metastatic behavior. While transcriptional plasticity is well recognized as a driver of melanoma state transitions, mechanisms linking chromatin-associated regulators to extracellular matrix remodeling remain poorly understood. Our data position HSPGs as key effectors of this process and indicate that EZH2 modulates HS biosynthesis and remodeling independently of its canonical PRC2-associated chromatin regulatory activity. Through integrated bioinformatic, genomic, and functional approaches, we demonstrate that EZH2 exerts dual and mechanistically distinct control over HS-modifying sulfatases through canonical PRC2-dependent repression of *SULF2* via H3K27 trimethylation and non-canonical, methyltransferase-independent activation of *SULF1* through a TRIM28-dependent mechanism. These opposing regulatory programs converge to alter HS sulfation patterning at the tumor cell surface, thereby modulating ligand binding and likely contributing to enhanced invasive and metastatic potential. Importantly, genetic disruption of *SULF1* impeded metastatic progression in vivo, highlighting extracellular HS remodeling as a key downstream effector of melanoma malignancy.

A central finding of this study is the mechanistic divergence by which EZH2 regulates the two extracellular HS 6-*O*-endosulfatases, SULF1 and SULF2. The expression of both SULF1 and SULF2 is dysregulated in multiple malignancies and has been associated with poor survival outcomes, with studies reporting both tumor-suppressive and oncogenic roles depending on cancer type and disease context^9^. These enzymes are often considered functionally redundant due to their shared ability to remove 6-*O* sulfate groups from extracellular HS chains^49, 50^. However, studies have reported context-dependent and sometimes opposing roles for SULF1 and SULF2 across cell types and cancer subclasses^51, 52, 53, 54, 55^. For example, SULF1 has been shown to play a regulatory role in cancer cell signaling, where it can be provided by cancer-associated fibroblasts to support metastasis and drug resistance in gastric cancer^56^. Conversely, SULF2 can be supplied by tumor cells to modulate key tumor intrinsic signaling pathways^53, 57, 58^. In chondrosarcoma, EZH2 represses SULF1 expression, and SULF1 functions as a tumor suppressor by attenuating cMET signaling^59^. In the current study, our data demonstrate that in melanoma, EZH2 activates SULF1 expression and alters HS 6-*O* sulfation through a non-canonical mechanism, and SULF1 acts as a pro-metastatic factor. These opposing regulatory roles for EZH2 in different cancers underscore that the functional consequences of SULF1 expression depend on tissue-specific signaling networks, HS-ligand interactions, and the epigenetic landscape of the tumor. The current results also expand upon our prior work, which reported that *SULF1* expression is suppressed by KDM2B^18^, a histone demethylase and PRC1 complex member, suggest divergent regulatory roles for PRC factors in melanoma cells. Together, our collective findings demonstrate that SULF1 and SULF2 are embedded within distinct epigenetic circuits, despite catalyzing similar biochemical reactions, and highlight how differential chromatin regulation can generate functional divergence among HS-modifying enzymes.

Mechanistically, we identify TRIM28 as a critical partner in non-canonical EZH2-mediated activation of SULF1 and other ECM- and proteoglycan-related pathways. Our study extends prior work that identified an EZH2/TRIM28 complex in MCF7 breast cancer cells and demonstrated that EZH2 and TRIM28 interact together with SWI/SNF co-factors to activate gene expression^44^. Our proteomic analysis of the EZH2 interactome in A375 cells revealed EZH2/TRIM28 interactions without detection of SWI/SNF components (Fig. 4A), suggesting a divergent mechanism in melanoma. These findings expand the functional repertoire of EZH2 beyond transcriptional repression and support a model in which EZH2 can act as a context-dependent transcriptional activator through protein-protein interactions. Importantly, this non-canonical function would not be fully suppressed by catalytic EZH2 inhibitors, providing a potential explanation for discrepancies between genetic and pharmacologic targeting of EZH2 observed in some cancer contexts^60, 61, 62^. Future studies will be required to determine whether EZH2/TRIM28 regulates *SULF1* through enhancer engagement, chromatin looping, or recruitment of additional transcriptional co-activators.

Functionally, we show that depletion of EZH2, TRIM28, or SULF1 impairs migration and invasion *in vitro*, and that SULF1 suppression alone reduces metastasis *in vivo* without markedly affecting primary tumor growth. These findings support a model in which SULF1 promotes the transition from proliferative to invasive states in melanoma, consistent with the role of HS-modifying enzymes in remodeling the tumor microenvironment. In fact, a recent study revealed that SULF1 overexpression in human melanoma cells leads to reduced tumor growth in xenografts via altered downstream AKT signaling^63^. Additionally, our data are consistent with prior studies reporting that targeting EZH2 or TRIM28 similarly impair cellular migration/invasion^25, 64, 65^. A recent study also highlighted the role of TRIM28 in regulating melanoma plasticity, where it can activate pro-invasive genes that control the balance between invasiveness and growth of melanoma cells^65^. Intriguingly, our tumor histology data suggest that knockdown of SULF1 may negatively impact primary tumor growth, which may reflect altered tumor-stromal interactions or vascular support. We speculate that enhanced HS sulfation may influence growth factor availability or cell-matrix interactions, potentially contributing to necrosis and reduced metastatic dissemination. Additionally, SULF1 depletion alone in the tumor cells resulted in reduced metastatic seeding, pointing to its intrinsic role in tumor migration, invasion, and metastasis. Although these xenograft studies were performed in immunocompromised mice and therefore do not capture immune contributions to metastasis, this model allows isolation of tumor-intrinsic effects of HSPG remodeling on melanoma progression. Forthcoming studies will address how EZH2/SULF1-mediated HS remodeling interfaces with immune regulation in melanoma.

In summary, this study defines the EZH2/TRIM28/SULF1 axis as a previously unrecognized mechanism linking epigenetic regulation to HS remodeling and melanoma metastasis. By uncovering a non-canonical function of EZH2 that shapes the extracellular matrix and tumor microenvironment, our findings broaden the conceptual framework for Polycomb protein function in cancer and highlight tumor-intrinsic mechanisms that govern the transition from primary growth to metastatic dissemination. While our mechanistic and functional analyses were performed primarily in BRAF-mutant melanoma models, which represent the most prevalent clinical subtype, future studies will be required to determine whether this regulatory axis similarly operates across additional melanoma genotypes and other tumor contexts. Importantly, SULF1 is an extracellular enzyme, rendering it more accessible to therapeutic intervention than nuclear epigenetic regulators. Targeting SULF1 or HS remodeling pathways may therefore represent a strategy to selectively impair metastasis, potentially reducing the toxicity associated with broadly acting epigenetic therapies and improving treatment durability.

## MATERIALS AND METHODS

### Bioinformatic analysis of transcriptional and epigenetic regulators of heparan sulfate

Predicted transcriptional and epigenetic regulators of heparan sulfate (HS) biosynthesis and heparan sulfate proteoglycan (HSPG) expression were identified using the offline toolkit of BART (Binding Analysis for Regulation of Transcription v2.1, https://zanglab.github.io/bart/) and ChIP-Atlas v3.0 (https://chip-atlas.org/). A curated gene list comprising enzymes involved in HS biosynthesis and modification, as well as core HSPGs, was used as input for both analyses. For BART, analyses were performed using the human genome build hg38, with enrichment based on overlap between input genes and transcription factor binding profiles from publicly available ChIP-seq datasets and ranked by BART score and associated p-values. ChIP-Atlas enrichment was performed using the “Enrichment Analysis” function (hg38) and experiment type “ChIP: TFs and others” (“All cell types”, threshold of significance = 50). Factors were ranked by enrichment score and statistical significance using “Random permutation of dataset A” for dataset B (permutation time = x1). For each platform, the top 100 significantly enriched regulators, by p-value, were selected and intersected to identify shared predicted regulators of HS biosynthesis for downstream analysis. Functional annotation and gene set enrichment analysis of the top 100 hits from each bioinformatic analysis was carried out using Metascape (http://metascape.org/)^66^. Overlapping hits were analyzed using STRING-DB (https://string-db.org/).

### Cell culture

A375 (CRL-1619), SK-MEL-5 (HTB-70), and HEK293T (CRL-3216) cells were grown in Dulbecco’s Modified Eagle Medium (DMEM, Gibco) supplemented with 10% (v/v) FBS and 1% penicillin/streptomycin (PenStrep) at 37°C under an atmosphere of 5% CO_2_/95% air. All cell lines were passaged every 3-4 days and revived from liquid nitrogen after ≤10 passages. All transfected cell lines were cloned and stored under liquid nitrogen.

### Plasmid preparation and cell line generation

An sgRNA targeting *EZH2* (5’–ATTGCTGGCACCATCTGACG–3’) was cloned into pSpCas9(BB)-2A-Puro (PX459, a gift from Feng Zhang and purchased from Addgene #62988) following the published procedure^67^. A375 wild-type cells (200,000 cells/well) were plated into 6- well plates and transfected with the plasmid with Lipofectamine LTX to generate *EZH2^-/-^* knockout cells. The mixed population of cells was then serial diluted into 96-well plates to establish clonal cell lines. Knockout clones were validated by Sanger sequencing and by Western blot. shRNA constructs were obtained from Sigma targeting *EZH2* (5’–CCAACACAAGTCATCCCATTA–3’, # TRCN0000040075), *SULF1* (5’– GCGAGAATGGCTTGGATTAAT–3’, #TRCN0000373588), *TRIM28* (5’– CTGAGACCAAACCTGTGCTTA–3’, #TRCN0000018001), and scrambled shRNA (5’– CCTAAGGTTAAGTCGCCCTCG–3’, Addgene #136035). HEK293T cells were used to generate lentivirus for each shRNA construct by transfection with Fugene 6 (Promega), envelope plasmid (pMD2.g, a gift from Didier Trono and purchased from Addgene, #12259), packaging plasmid (psPAX2, a gift from Didier Trono and purchased from Addgene, #12260), and shRNA constructs (4µg of each plasmid). Lentiviral particles were collected and subsequently added to A375 or SKMEL5 wild-type cells. Transduced cells were selected by treated with puromycin (1 µg/mL) to generate shRNA-expressing cell lines. A375 shCTRL-Luc and shSULF1-Luc cells were generated by transducing A375 shCTRL and shSULF1 cells with pLenti PGK Blast V5-Luc plasmid (a gift from Eric Campeau & Paul Kaufman and purchased from Addgene, #19166), followed by selection with blasticidin (2 µg/mL) for 3 days.

### Western blotting

Total protein was isolated from cells using RIPA Lysis and Extraction Buffer (EMD Millipore) supplemented with a protease inhibitor cocktail (Roche). Histones were isolated from cells using TEB buffer (PBS supplemented with 0.5% Triton X-100 and protease inhibitors) and acid extracted using 0.2M HCl overnight at 4°C. Protein concentrations were quantified using BCA assay (Thermo Scientific). Equal amounts of protein were loaded and separated on SDS-PAGE gels (4-12% Bis-Tris, NuPAGE, Invitrogen), transferred onto nitrocellulose membranes (0.22µm pore size, for histones) or polyvinylidene difluoride (PVDF) membranes (0.45µm pore size, for all other samples), and blocked with Intercept Blocking Buffer (LI-COR Biosciences) or 5% milk in Tris-buffered saline and 0.1% Tween (TBS-T) for 1 hour at room temperature. Membranes were probed with primary antibodies against EZH2 (Cell Signaling Technology, #5246; 1:1000), EZH2 (Active Motif, #39076, 1:200), TRIM28 (Cell Signaling Technology, #85322; 1:1000), SUZ12 (Cell Signaling Technology, #3737; 1:1000), EED (Cell Signaling Technology, #85322; 1:1000), H3K27me3 (Cell Signaling Technology, #9733; 1:1000), Histone H3 (Cell Signaling Technology, #3638; 1:1000), or β-actin (Cell Signaling Technology, #3700; 1:5000) at 4°C overnight. After washing with TBST, mouse and rabbit primary antibodies were incubated with secondary Odyssey IR dye antibodies (anti-mouse 800, LI-COR Biosciences, #926-32212; donkey anti-rabbit 680, LI-COR Biosciences, #926-68073; 1:14,000) and visualized on an Odyssey infrared imaging system (LI-COR Biosciences). Uncropped western blot source images are included in the Supplementary Information (Suppl. Figs. 7-8).

### Rescue experiments and siRNA transfections

For EZH2 rescue experiments, we utilized a plasmid containing human *EZH2* gene (EZH2-WT) obtained from ABM (pLenti-GIII-CMV-EZH2-HA, Applied Biological Materials). The EZH2-H694A mutant plasmid was generated from the EZH2-WT vector by inducing a single amino acid substitution (H694A) using the Q5 Site-Directed Mutagenesis Kit (New England Biolabs). A375 wild-type or *EZH2^-/-^* knockout cells (2 × 10^5^ cells/well) were transfected with Lipofectamine LTX with Plus reagent (Invitrogen) and EZH2-WT or EZH2-H694A (100 ng each). Cells were incubated for 72 hours before performing downstream experiments.

For SULF1 rescue experiments, *EZH2^-/-^*cells (2 × 10^5^ cells/well) were transfected with Lipofectamine LTX with Plus reagent (Invitrogen) and a human SULF1 expression plasmid (gift from Steven Rosen and obtained from Addgene, # 24425). For knockdown experiments, A375 wild-type cells (2 × 10^5^ cells per well) were transfected with Lipofectamine LTX with Plus reagent (Invitrogen) and an siRNA targeting human *EED* (SASI_Hs02_00336316; Sigma), *SUZ12* (SASI_Hs02_00347682; Sigma), or a non-targeting negative control siRNA (#SIC001, Sigma) according to the manufacturer’s instructions. Cells were incubated with each mixture for 4 hrs, after which the media was replaced with DMEM (+ 10% FBS). FACS binding and qPCR experiments were performed 48 hrs post-transfection.

### Heparan sulfate isolation, purification and LC-MS disaccharide analysis

A375 wild-type or *EZH2^-/-^*cells (5×10^5^) were seeded in 10 cm plates and grown to ∼90% confluency. Cells were then washed with PBS, lifted with trypsin, and pelleted. The trypsin fraction containing released glycosaminoglycans was collected and treated with Pronase (0.5mg/mL, Sigma) overnight at 37°C. The product was filtered and passed through a DEAE-Sephacel column equilibrated with wash buffer (50 mM sodium acetate, 200 mM NaCl, 0.1% Triton X-100 (v/v), pH 6.0). The sample was subsequently passed through a PD-10 desalting column (Cytiva) and lyophilized overnight. For HS disaccharide analysis, purified GAGs were incubated with 2 mU each of heparin lyases I, II, and III (IBEX) for 16 hrs at 37°C in heparinase buffer (40mM sodium acetate, 3.3mM calcium acetate, pH 7.0). HS disaccharides were aniline-tagged and analyzed by RP-LC/MS on a LTQ XL Orbitrap mass spectrometer, as previously described^68^. Protein concentrations from cell pellets were quantified for each sample by BCA assay and used to determine total cell surface HS levels. All LC-MS disaccharide analysis results are included in Supplementary Tables 3-4 in the supplementary information.

### RNA extraction and quantitative PCR

Total RNA was isolated from cells using TRIzol (Invitrogen) and the Direct-zol RNA Miniprep kit (Zymo Research) according to manufacturer’s instructions. iScript™ cDNA Synthesis kit (Bio-Rad) was used to prepare cDNA following the manufacturer’s instructions. Quantitative PCR (qPCR) was performed with cDNA and PowerUp SYBR Green Master Mix (Applied Biosystems) following the manufacturer’s instructions. The expression level of *YWHAZ* was used to normalize target gene expression across samples. The primers used for gene expression analyses are listed in Supplementary Table 2.

### RNA sequencing and differential expression analysis

For transcriptome analysis, total RNA from shCTRL, shEZH2, and shTRIM28 knockdown cell lines were used for library preparation and next-generation sequencing (Azenta). Raw sequencing reads were trimmed using Trimmomatic v.0.36 and mapped to the human reference genome (GRCh38) using the STAR aligner v.2.5.2b. The Subread package v.1.5.2 was used to calculate the unique gene hit counts by using featureCounts. Differential gene expression analysis was performed using DESeq2, considering genes with an FDR p-value ≤ 0.01 and a fold change of ± 2 as differentially expressed. Log_2_ fold changes and p-values were generated using the Wald test. Functional annotation and gene set enrichment analysis of the top differentially expressed genes were performed using Metascape (http://metascape.org/)^66^.

### Protein biotinylation

Heparin-Sepharose (100 μl, Cytiva) was pre-equilibrated with PBS (Gibco) and then loaded with human FGF1 (Peprotech, #100-17A) or human FGF2 (Peprotech, #100-18B) dissolved in PBS, as previously described^34^. The flow-through was collected and reloaded onto the column twice to ensure complete binding. After washing twice with PBS, 0.6 mg/mL of Sulfo-NHS-LC-biotin (Thermo Fisher) in PBS was applied to the column and incubated for 1 hour at room temperature. Each column was washed three times with PBS, then bound biotinylated protein was eluted with 0.4 mL of PBS buffer containing an additional 2 M NaCl. All biotinylated proteins were stored at −80°C before use.

### Flow cytometry

A375 and SKMEL5 cells grown in monolayer were lifted using 10 mM EDTA in PBS and incubated in suspension for 30 min (1 hr for anti-thrombin) on ice in PBS (+ 0.1% BSA) with 80 nM biotin-FGF1, 2.5 nM biotin-FGF2, 500 nM anti-thrombin (Aniara Diagnostica, #APP004B), 0.5 µg/mL mAb 10E4 (AMSBio, #370255-1, Clone F58-10E4, 1:2000), or 1 µg/mL mAb 3G10 (AMSBio, #370260-S, clone F69-3G10, 1:1000). Heparin lyase pre-treatment was performed prior to 3G10 binding by incubating lifted cells with 5 mU/mL of heparin lyases I, II, and III (IBEX) at 37°C for 30min in PBS (+ 0.1% BSA). Biotinylated proteins were detected by Streptavidin-Cy5 (Invitrogen, 1:1000). Bound anti-thrombin was detected by 2 µg/mL goat IgG anti-AT-III pAb (R&D Systems, AF1267, 1:100), followed by 2.5 µg/mL donkey anti-goat IgG conjugated with AlexaFluor 488 (Invitrogen, #A-11055). 10E4 was detected by 2 µg/mL anti-mouse IgM AlexaFluor 647 (Invitrogen, #A-21238, 1:1000). Bound 3G10 was detected by 2 µg/mL anti-mouse IgG AlexaFluor 488 (Invitrogen, #A-11001, 1:1000). Flow cytometry was performed using a CytoFLEX S instrument (Beckman Coulter; ≥10,000 events/sample). Live cells were gated based on forward and side scattering. The extent of protein binding was quantified using the geometric mean of the fluorescence intensity. Unstained negative controls were included in each experiment. Data were plotted and analyzed in GraphPad Prism v8.0.

### Radiolabeling and size exclusion chromatography

A375 wild-type and *EZH2^-/-^* cells (2×10^5^) were plated in 6-well plates, let adhere overnight, then incubated in F12 medium supplemented with 10% dialyzed FBS for 30min. Cells were radiolabeled with 20 µCi/mL [^35^S]sulfate in F12 (+ 10% dialyzed FBS) for 24 hrs at 37°C. HS chains were purified as described above and analyzed by size exclusion chromatography (Sepharose CL-6B column 1.7 cm x 80 cm; 50mM sodium acetate, 0.2M NaCl, pH 6.0). HS average molecular mass was determined based on previous size determinations using a Sepharose 6B column^69^.

### Drug treatments

All drugs were dissolved in DMSO. 2×10^5^ cells were seeded into 6-well plates and treated with media containing DMSO (vehicle) as a negative control or EZH2 inhibitors EPZ-6438 (2 µM), GSK126 (2 µM), and DZNep (1 µM) for 48 hours. Total RNA was then purified from cells and analyzed using qPCR, as detailed above. Total proteins and histones were isolated as detailed above for Western blot analyses.

### Co-immunoprecipitation and LC-MS/MS proteomics

A375 wild-type cells (5×10^6^) cells were seeded in 15 cm plates. Cells were harvested and resuspended in IP Lysis Buffer (Thermo Scientific) supplemented with protease inhibitors (Sigma-Aldrich). Protein lysates (2.5 mg/IP) were then applied to protein G Dynabeads (Thermo Scientific) conjugated to antibodies (2.5 µg per sample) targeting EZH2 (Active Motif, #39076), TRIM28 (Abcam, #ab10483), or IgG isotype control (Cell Signaling Technology, #2729) and incubated overnight at 4°C, with rotating. Protein G Dynabead conjugates were recovered from solution using a magnetic rack and subsequently washed three times with IP Antibody Washing Buffer (with 0.1% Tween-20) using the Dynabeads Immunoprecipitation Kit (Thermo Scientific). Beads were then washed three times with IP Antibody Washing Buffer (without Tween-20) and proteins were eluted from beads using elution buffer (50mM triethylammonium bicarbonate, 5% SDS) and boiled at 95°C for 10 minutes. Eluted protein samples were then reduced by incubating with 120 mM tris (2-carboxyethyl) phosphine (TCEP) at 56°C for 15 minutes and alkylated by incubation with 500 mM methyl methanethiosulfonate (MMTS) at room temperature for 10 minutes. After alkylation, the proteins were washed over suspension trapping columns (ProtiFi) and then digested using trypsin/LysC mix (Promega) at 37°C overnight.

The resulting peptides were separated on an Acclaim™ PepMap™ 100 C18 column (75 µm x 15 cm) and eluted into the nano-electrospray ion source of an Orbitrap Eclipse™ Tribrid™ mass spectrometer (Thermo Scientific) at a flow rate of 200 nL/min. The elution gradient consists of 1-40% acetonitrile in 0.1% formic acid over 320 minutes followed by 10 minutes of 80% acetonitrile in 0.1% formic acid. The spray voltage was set to 2.2 kV and the temperature of the heated capillary was set to 275 °C. Full MS scans were acquired from m/z 300 to 2000 at 60k resolution, and MS/MS scans following collision-induced dissociation (CID) at 38% collision energy were collected in the ion trap. The spectra were analyzed using SEQUEST (Proteome Discoverer 2.5, Thermo Fisher Scientific) with mass tolerance set as 20 ppm for precursors and 0.5 Da for fragments. The search output was filtered to reach a 1% false discovery rate at the protein level and 10% at the peptide level. Quantitation was performed by calculating spectral counts for each protein.

### Chromatin immunoprecipitation (ChIP)

For EED and H3K27me3 IP, ∼6×10^6^ cells were initially crosslinked with 1% formaldehyde diluted in PBS (Sigma Aldrich) at RT for 10 minutes and then quenched with 2.625 M glycine for 3 minutes. The cells were then washed twice with PBS and isolated into Eppendorf LoBind tubes. Lastly, the cells were pelleted and the supernatant was discarded. Cell pellets were then snap-frozen with liquid nitrogen to be used for library preparation. Cells pellets were resuspended in 300 µl ice-cold LB3 lysis buffer (10 mM Tris-HCl, pH 8.0, 100 mM NaCl, 1 mM EDTA, 0.5 mM EGTA, 0.1% Na-Deoxycholate, 0.5% N-lauroylsarcosine, 1x protease inhibitor cocktail [PIC]), and the lysed samples were sonicated using the Bioruptor Pico device with the following settings: 10 cycles, 10s ON, 50s OFF, on medium intensity. Next, lysed and sonicated samples were diluted using 30 µl of 10% Triton X-100 and centrifuged for 10 minutes at maximum speed. 1% of the volume of supernatant was set aside to be used as input sample. The remaining supernatant was transferred to new Eppendorf tubes and incubated overnight with 2 µg of antibody targeting EED (Abcam, #AB240650) or H3K27me3 (Active Motif, #39155) and 20 µl of Protein A Dynabeads on a rotator at 4°C overnight. The samples were placed under a magnetic field, and the supernatant is discarded. The beads were then washed 3x with Wash Buffer I (20 mM Tris-HCl/pH 7.4@20°C, 150 mM NaCl, 0.1% SDS, 1% Triton X-100, 2 mM EDTA, 1x PIC), 3x WB III (250 mM LiCl, 1% Triton X-100, 0.7% DOC, 10 mM Tris-HCl/pH 7.4@20°C, 1 mM EDTA, 1x PIC), 2x TET (0.2% Tween 20/TE, 1x PIC) for 30 seconds and 1x TE-NaCl (50mM NaCl, 1x PIC). Lastly, the beads were resuspended in 25 ul of TT (10mM Tris pH 8.0, 0.05% Tween-20) for on-bead library preparation.

For TRIM28 IP, ∼6×10^6^ cells were first crosslinked using 2 mM of disuccinimidyl glutarate (Pierce Cat # 20593) diluted in PBS for 30 minutes at RT, formaldehyde was added to a final concentration of 1% with an additional incubation of 10 minutes at RT. The following steps were identical to EED/H3K27me3 ChIP. TRIM28 IP was performed with 2 µg of antibody (Invitrogen, #20C1). Libraries from 2 biological replicates were prepared for each condition, including an input sample, which corresponded to single-crosslinked, fragmented cells where no antibody was used for enrichment. Libraries were prepared with KAPA HyperPrep library preparation kit (Roche) using half reactions. IDT adapters were used for pooling samples for next-gen sequencing. Chromatin was eluted and crosslinks were reversed, and the DNA in the eluate was amplified for 14 cycles. Lastly, double-sided size selection was performed on the DNA libraries to ensure the enrichment of desirably sized fragments. The first selection was performed using 0.56x KAPA beads and the second with 0.8x. The resulting libraries were paired-end sequenced (2×150bp) on the Illumina NovaSeq X Plus platform by the Core Facility Genomics (CFG) at Amsterdam UMC.

### ChIP sequencing analysis

The fastq files of the sequenced libraries were obtained by the CFG and quality control was performed on the raw sequencing data using the FastQC tool. After quality of samples was confirmed, reads were trimmed and Illumina universal adapter sequences were removed using Trimmomatic v0.39 in paired-end mode using the following parameters: ILLUMINACLIP:TruSeq3-PE-2.fa:2:30:10 LEADING:3 TRAILING:3 SLIDINGWINDOW:4:15 MINLEN:36. Unpaired reads were discarded from subsequent analysis, while successful read pairs were aligned to the human genome assembly hg38 using the alignment program HISAT2 2.1.0 using the parameters “--mm --add-chrname --new-summary --no-spliced-alignment”.

Output sam files were converted to bam format, and secondary, supplementary and reads with a mapping quality (MAPQ) lower than 20 were discared using samtools 1.17. Duplicate reads originating from a single fragment of DNA were removed using Picard’s MarkDuplicates (version 2.9.0-1-gf5b9f50-SNAPSHOT). BigWig files were generated using deeptools’ bamCoverage and read coverage was normalized using the RPGC method (reads per genomic content [1x normalization]). Peak calling was performed using MACS3 with the input sample used as control. Peaks for H3K27me3 and EED were called in “broad” peak mode using a broad cutoff of 1e-02 and a q-value of 1e-03. Peaks for TRIM28 were called in “narrow” peak mode a q-value of 1e-03. A “consensus” peak set for each IP was determined by selecting the peaks that overlap in the two replicates by at least 100 base pairs (broad) or 50 base pairs (narrow). Peaks were annotated using ChIPseeker (1.46.1) in R. Line plots depicting the signal of EED, TRIM28 and H3K27me3 around gene bodies were generated using deepTools’ computeMatrix scale-regions.

### Cell viability, migration, and invasion assays

For growth curves, cell lines were plated in 96-well plates at a density of 3×10^3^ cells per well in DMEM media containing 2% FBS and 1% PenStrep. Cell Titer Blue (Promega) was used to measure cell viability every 24 hours for 5 days, following the manufacturer’s instructions.

For migration assays, Transwell™ inserts (Corning) with 8 µm pore sizes were used. Cell lines were serum-starved overnight in DMEM supplemented with 1% PenStrep. Cells were then lifted with trypsin, resuspended in low-serum media (DMEM +0.2% FBS, 1% PenStrep), and seeded into Transwell™ inserts at a density of 2.5×10^4^ cells per insert. DMEM containing 10% FBS (+ 1% PenStrep) was added to the lower chamber, and cells were incubated at 37°C under an atmosphere of 5% CO2/95% air for 48 hours prior to analysis.

For invasion assays, Matrigel™ GFR invasion chambers with 8µm pore sizes were used. Cells were lifted with trypsin, resuspended in low-serum media (DMEM +0.2% FBS, 1% PenStrep), then seeded into Matrigel-coated chambers at a density of 5.0×10^4^ for A375 cells and 1×10^5^ for SKMEL5 cells per insert. DMEM containing 10% FBS (+ 1% PenStrep) was added to the lower chamber, and cells were incubated at 37°C under an atmosphere of 5% CO2/95% air for 36 hours prior to analysis.

At the endpoint, each insert for migration and invasion assays was fixed with 4% paraformaldehyde then permeabilized with 100% methanol and stained with 0.2% crystal violet (w/v in 10% ethanol). Each insert was imaged by microscopy using 10X magnification. For quantification, crystal violet was eluted using 33% acetic acid solution and absorbance was measured at 590 nm using a plate reader.

### Luminescence assays

The luminescence signal of A375 shCTRL-Luc and shSULF1-Luc cells was quantified by incubating cells with 750 µg/mL D-luciferin (GoldBio) for 5 minutes at room temperature followed by measuring luminescence using a plate reader. Cells lacking luciferase expression were used as a negative control.

### Orthotopic In vivo Tumor Studies

All animal experiments were conducted in accordance with a protocol approved by Virginia Commonwealth University (VCU) Institutional Aimal Care and Use Committee and conformed to NIH guidelines. Male and female NSG (NOD.Cg-Prkdcscid Il2rgtm1Wjl/SzJ) mice, aged 6 weeks, were obtained from VCU Cancer Mouse Models Core (CMMC), which maintains colonies derived from initial breeders purchased from The Jackson Laboratory (JAX). Cells were confirmed negative for *Corynebacterium bovis* and *Mycoplasma* contamination prior to experiments.

Mice were injected intradermally into both hind flanks with 1×10^6^ A375 shCTRL-Luc or shSULF1-Luc cells diluted in PBS. Body weight was monitored twice weekly. Tumor volume (V) was calculated with the formula: V (mm^3^) = 0.5 x L x W^2^ where L= length and W=width. Bioluminescent imaging was performed with the IVIS Lumina S5 (Revvity). Total flux (photons/sec) and tumor volume were recorded in Studylog lab management software.

Mice were euthanized when the total primary tumor volume reached ∼2000 mm^3^. At endpoint, mice were imaged live, euthanized, and tumors were harvested and weighed. Organs including lungs, brain, liver, and bones were collected to assess metastases by ex vivo IVIS imaging. Each organ was immersed for 5 min in luciferin/PBS (0.3 mg/ml) prior to imaging.

### Histopathology

Tissues and tumors were fixed in 10% neutral buffered formalin for at least 3 days, then paraffin embedded and sectioned at 5 µm. Hematoxylin and eosin (H&E) staining was performed in the VCU Tissue and Data, Acquisition and Analysis Core. H&E staining was automated using the Coverslipper (Agilent Technologies). Slides were scanned using the PhenoImager HT (Akoya Biosciences) at 20X magnification. Visualization was performed with Phenochart Whole Slide Contextual Viewer, and necrotic areas were quantified using QuPath software (v0.6.0, https://qupath.github.io/), with confirmation by Dr. Mochel (board certified pathologist).

### TCGA clinical data analysis

RNA-seq data showing gene expression profiles of primary and metastatic tumors from patients were obtained from the TCGA-SKCM (skin cutaneous melanoma) project dataset. Transcripts per million (TPM) expression data were extracted and plotted to compare gene expression quantification of *EZH2, TRIM28,* and *SULF1* between primary and metastatic samples.

### Statistics and reproducibility

Statistical tests, sample sizes, and numbers of biological replicates are indicated in figure legends. Statistical significance was indicated by the following: *p<0.05, **p<0.01, ***p<0.001. All tests were two-tailed and performed using GraphPad Prism v.8.0 or v10.6 software. Measurements were taken from distinct biological samples, and the number of biological replicates is indicated in figure legends. Error bars in plotting total tumor volume over time and histological analyses represent mean ± standard error of the mean. All other error bars represent mean ± standard deviation. Western blots, migration and invasion assays were performed three times independently and representative images are shown in the figures.

## Data availability

Raw data for LC-MS analysis of GAGs are available at GlycoPOST^70^ under project ID GPST000658. All mass spectrometry data has been deposited to ProteomeXchange Consortium via the MassIVE partner repository (https://massive.ucsd.edu/ProteoSAFe/static/massive.jsp) with the dataset identifier MSV000098756. Raw sequencing reads and results of the RNA and ChIP sequencing analyses will be made available at NCBI Gene Expression Omnibus upon publication. Any data supporting the analyses in the manuscript are available from the corresponding author upon reasonable request.

## Supplementary Information

Supplementary tables, figures, and source data are provided in the Supporting Information.

## Acknowledgements

We thank IBEX Technologies for their in-kind donation of heparin lyase enzymes. We also thank Dr. Biswa Choudhury and the GlycoAnalytics Core Facility at University of California, San Diego for help with analytical experiments, and Dr. Kosuke Funato (UGA) for providing reagents for the luciferase assays. This work was supported by UGA Startup funds and NIH grants to R.J.W. (NIGMS R35GM150736), the NSF BioFoundry: Glycoscience Research, Education, and Training grant (BioF:GREAT, 240020, to L.W.), and an ERC Starting Grant from the European Union (ERC, CytoMAC, 101076170) to M.A.H. N.G.P. was partially supported by a NIGMS T32 training grant (GM107004) and ARCS Foundation Scholarship. Services and products in support of the research project were generated by the Virginia Commonwealth University Cancer Mouse Models Core Laboratory and the Tissue and Data Acquisition and Analysis Core Laboratory, supported, in part, with funding to the Massey Cancer Center from NIH-NCI Cancer Center Support Grant P30 CA016059.

## Author Contributions

Unless otherwise noted, N.G.P. performed the experimental work and analyzed the data. A.D., M.B., and M.A.H. performed ChIP-seq experiments and analyzed the data. F.V. and E.S. characterized knockout cell lines and performed immunoblotting and qPCR experiments. A.B. generated, processed, and analyzed the LC-MS GAG compositional data. J.C.M. performed radiolabeling experiments and size exclusion analyses. X.D., P.Z. and L.W. generated, processed, and analyzed the proteomics data. B.H. and J.E.K. performed mouse tumor studies. M.C.M. reviewed and provided expert analysis of histology data. N.G.P. and R.J.W. wrote the paper, with input from all listed co-authors.

---

**Supplementary Figure 1.**
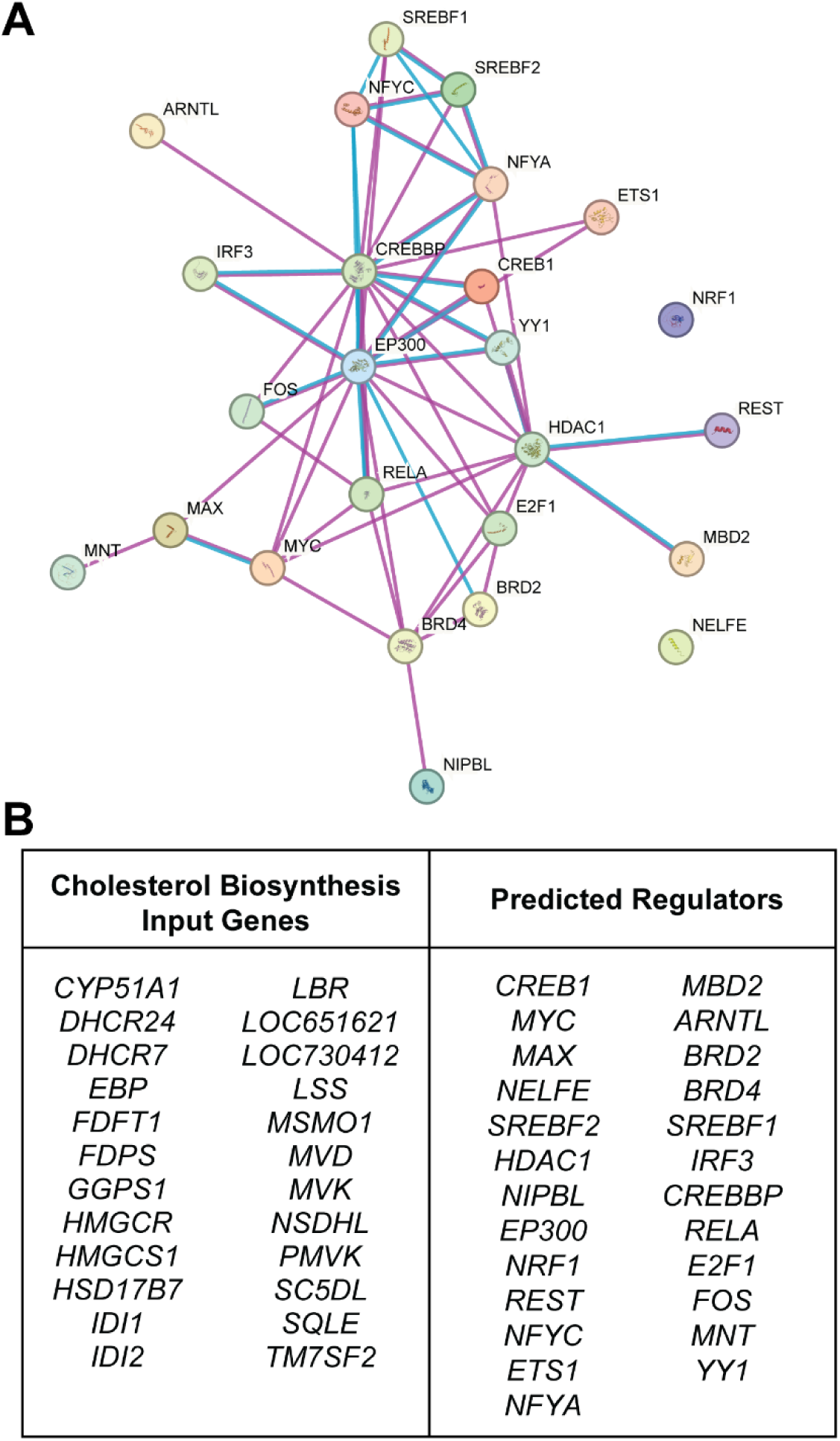
BART and ChIP-Atlas predict known regulators of cholesterol biosynthesis. **(A)** STRING-DB analysis showing interactome of overlapping predicted regulators of cholesterol biosynthesis generated from BART and ChIP-Atlas analyses. **(B)** Table displaying BART and ChIP-Atlas input gene list for cholesterol biosynthesis (left) and prediction results for potential regulatory factors enriched across both bioinformatic tools.

**Supplementary Figure 2.**
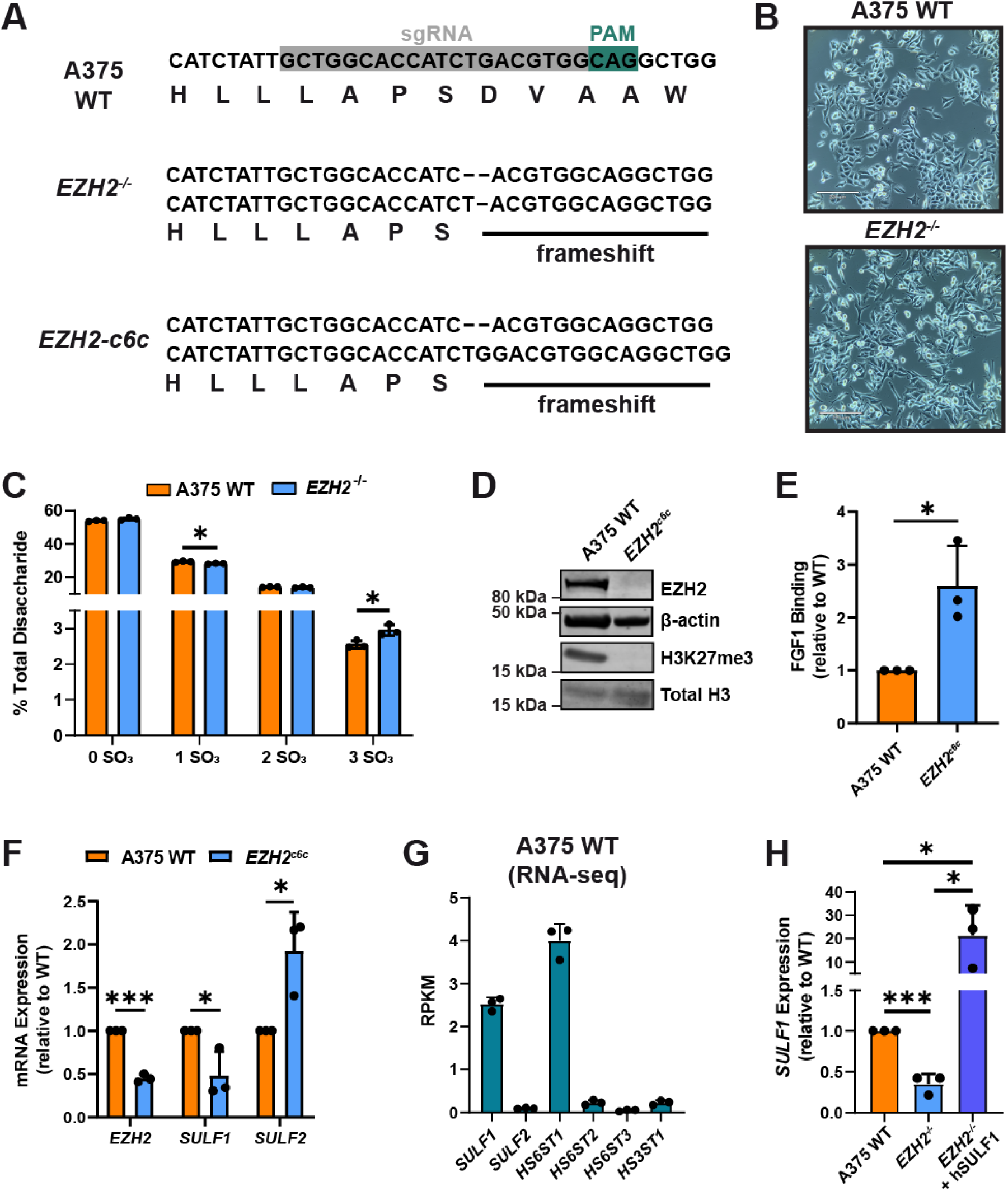
Characterization of A375 *EZH2* knockout cell lines. **(A)** Sanger sequencing of two A375 *EZH2* knockout clones. **(B)** Microscopy image of A375 wild-type and *EZH2^-/-^* cells. Scale bar: 220 µm. **(C)** LC-MS analysis of number of sulfates per disaccharide for cell surface HS isolated from A375 wild-type and *EZH2^-/-^* cells. **(D)** Western blot of EZH2 and H3K27me3 levels in A375 wild-type and a second *EZH2* knockout cell line (*EZH2*^c6c^). **(E)** Flow cytometry analysis of FGF1 binding in A375 wild-type and *EZH2*^c6c^ cells. **(F)** qPCR analysis of *EZH2*, *SULF1*, and *SULF2* mRNA expression in A375 wild-type and *EZH2*^c6c^ cells. **(G)** Basal gene expression (RPKM) of genes encoding a subset of HS biosynthetic enzymes. Data was extracted from a published study (GSE224599)^19^. **(H)** qPCR of *SULF1* expression in A375 wild-type, *EZH2^-/-^*, and *EZH2^-/-^* cells transfected with a human SULF1 expression plasmid. Data are presented as mean ± SD (n = 3 independent biological replicates), ***p<0.001, **p<0.01, *p<0.05 by two-sided t-test.

**Supplementary Figure 3.**
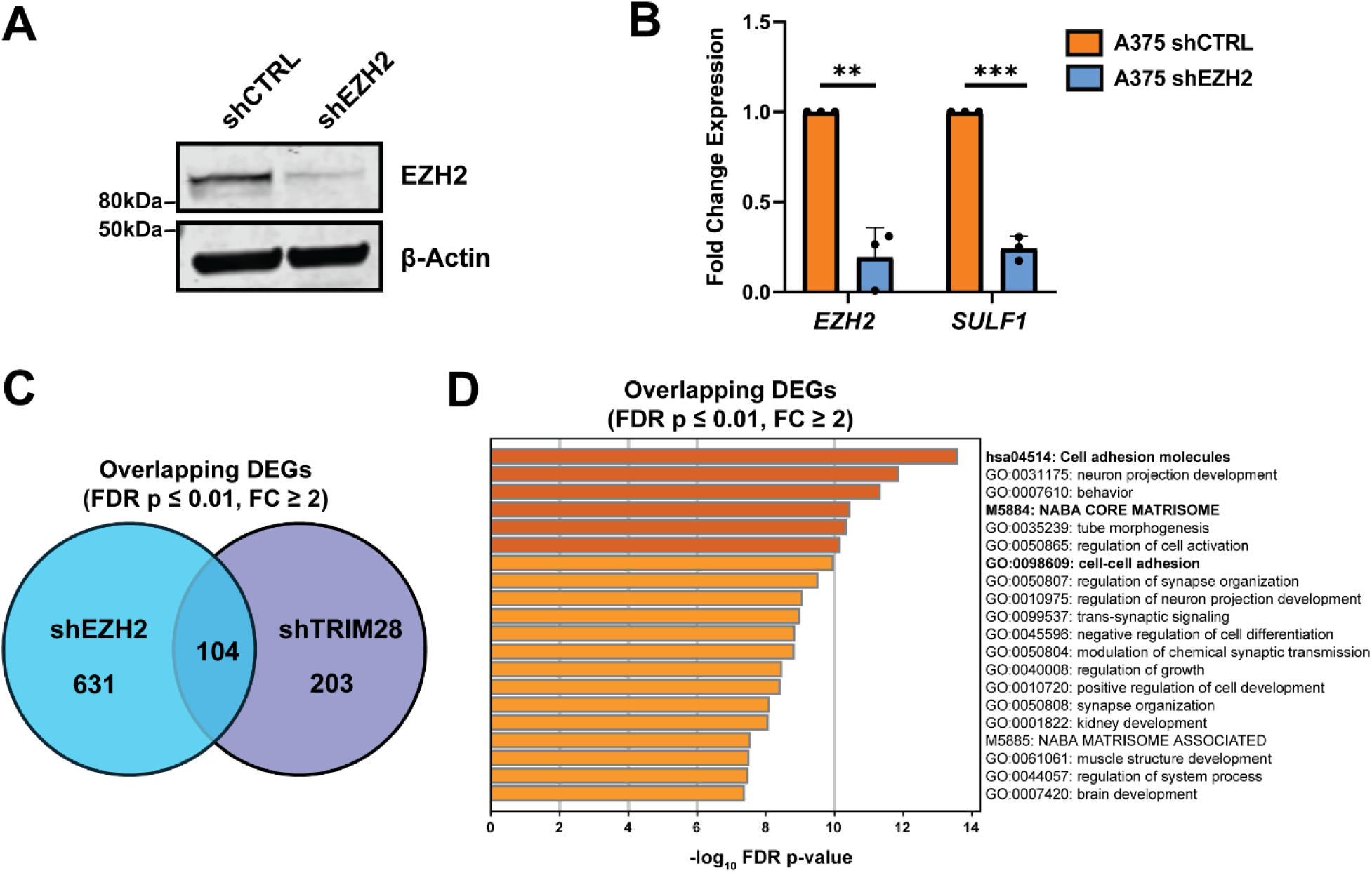
shRNA validation experiments and RNA-seq pathway analysis. **(A)** Western blot showing depleted EZH2 protein in shEZH2 cells. **(B)** qPCR analysis of *SULF1* mRNA expression upon stable knockdown of *EZH2*. **(C)** Venn diagram showing overlap of all DEGs in shEZH2 and shTRIM28 RNA-seq datasets (FDR p-value ≤ 0.01, FC ≥ 2). **(D)** Pathway enrichment analysis of all overlapping DEGs between shEZH2 and shTRIM28 populations (FDR p-value ≤ 0.01, FC ≥ 2). Data are presented as mean ± SD (n = 3 independent biological replicates, n = 4 independent biological replicates for RNA-seq experiments), ***p<0.001, **p<0.01, *p<0.05 by two-sided t-test.

**Supplementary Figure 4.**
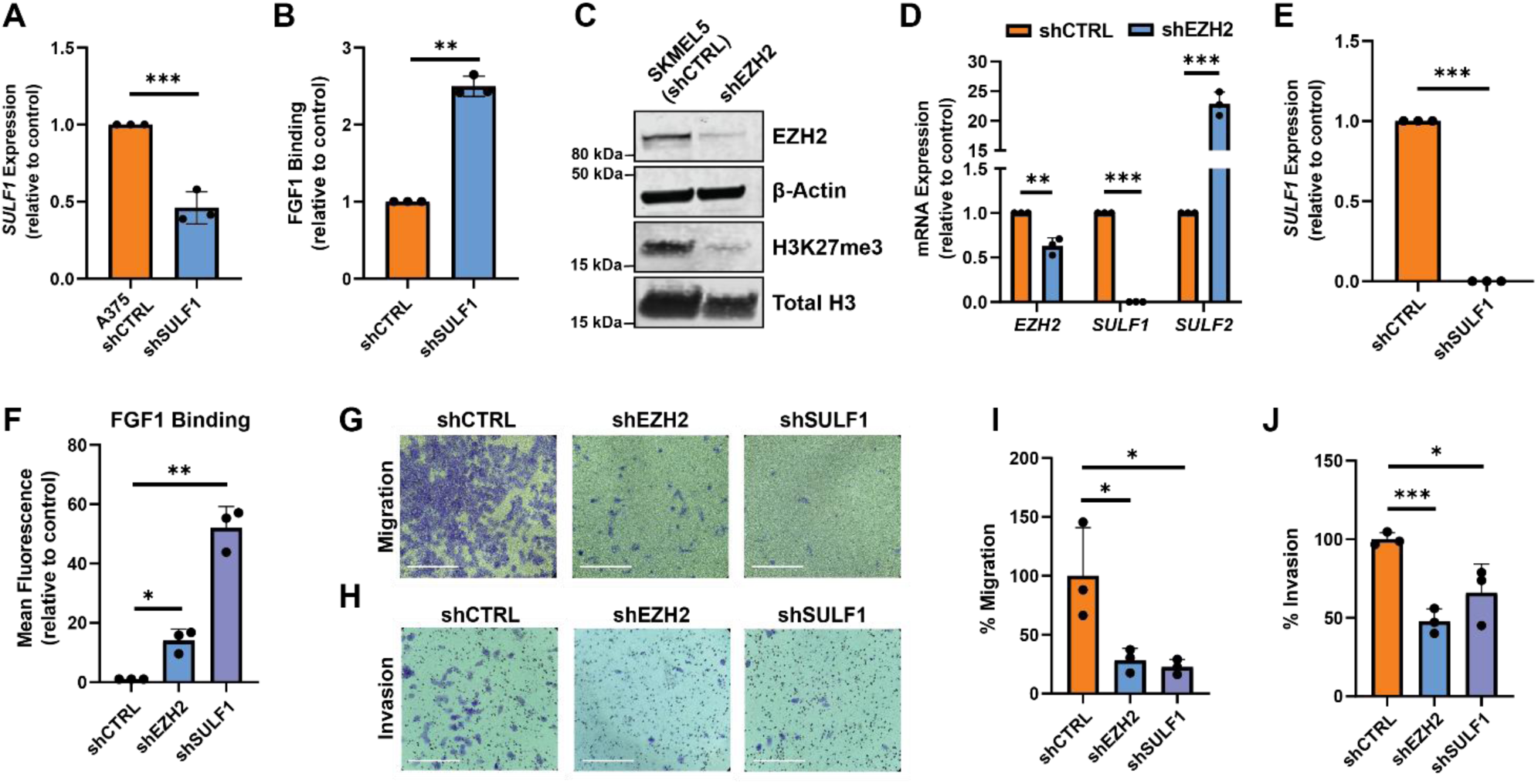
Characterization of A375 and SKMEL5 shRNA knockdown cells. **(A)** qPCR analysis showing reduced *SULF1* mRNA expression upon shRNA-mediated *SULF1* knockdown in A375 cells. **(B)** Flow cytometry analysis of cell surface binding of FGF1 in A375 shSULF1 compared to shCTRL cells. **(C)** Western blot showing depleted EZH2 protein and reduced global H3K27me3 levels in SKMEL5 EZH2 knockdown cells (shEZH2) relative to SKMEL5 wild-type control cells (shCTRL). **(D)** qPCR analysis showing altered *SULF1* and *SULF2* mRNA expression upon shRNA-mediated *EZH2* knockdown in SKMEL5 cells. **(E)** qPCR validating shRNA-mediated knockdown of *SULF1* expression in SKMEL5 cells. **(F)** Flow cytometry analysis of cell surface binding of FGF1 in SKMEL5 shEZH2 and shSULF1 cell lines compared to shCTRL cells. **(G)** Representative images of migrated cells stained with crystal violet (48-hour incubation). Scale bar: 360 µm. **(H)** Representative images of Matrigel invading cells stained with crystal violet (36-hour incubation). Scale bar: 360 µm. **(I)** Quantification of cell migration assays by measuring absorbance (590 nm) of crystal violet eluted from transwell inserts. Data was normalized to shCTRL cells. **(J)** Quantification of cell invasion assays by measuring absorbance (590 nm) of crystal violet eluted from Matrigel inserts. Data was normalized to shCTRL cells. Data are presented as mean ± SD (n = 3 independent biological replicates), ***p<0.001, **p<0.01, *p<0.05 by two-sided t-test.

**Supplementary Figure 5.**
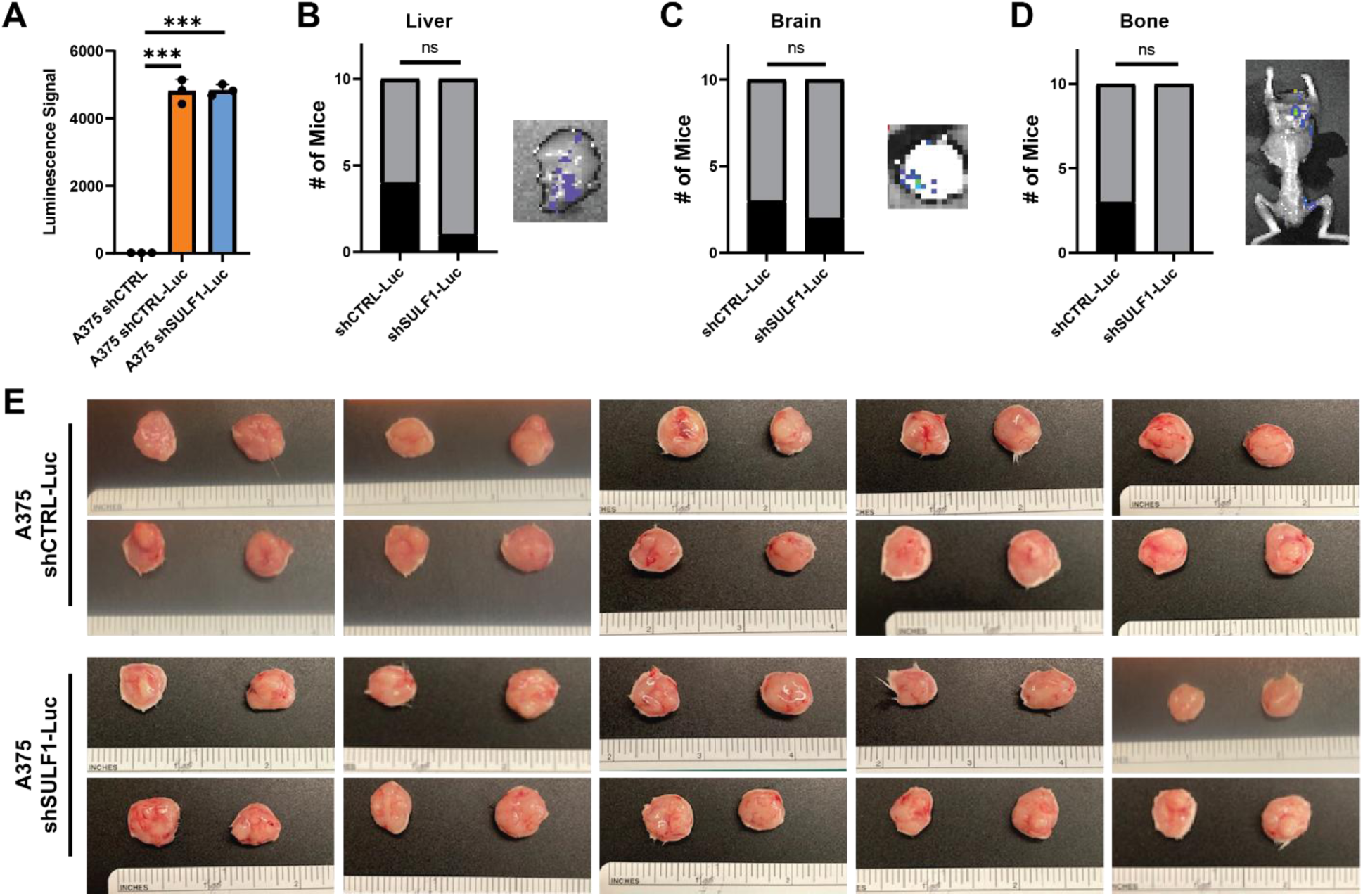
Luciferase assays, metastases, and endpoint *ex vivo* images of primary tumors. **(A)** Quantification of luminescence for A375 shCTRL-Luc and shSULF1-Luc cell lines. Contingency analysis showing the number of mice with detectable metastases, as assessed by ex vivo bioluminescence imaging, in **(B)** liver, **(C)** brain, and **(D)** bone (n = 10 mice per group). **(E)** Images taken of each tumor excised from mice once total tumor volume reached ∼2,000 mm^3^. Two primary tumors were harvested from each mouse. Data are presented as mean ± SD, ***p<0.001, **p<0.01, *p<0.05, unless otherwise stated above. Statistical significance was determined using two-sided tests as indicated (luciferase assay: two-sided t-test; metastasis incidence: chi-squared test).

