## Supplementary Information for "A non-canonical EZH2/TRIM28 epigenetic axis drives heparan sulfate remodeling and melanoma metastasis"

**This PDF file includes:**

- Supplementary Tables 2-4
- Supplementary Figures 6-8

**Supplementary Table 2:**
Primer sequences used for quantitative PCR and ChIP-qPCR analyses

| Gene | Forward Primer Sequence (5’→3’) | Reverse Primer Sequence (3’→5’) |
|---|---|---|
| *YWHAZ* | CCTGCATGAAGTCTGTAACTGAG | GACCTACGGGCTCCTACAACA |
| *EZH2* | AATCAGATGCGACTGAGA | GCTGTATCCTTCGCTGTTTCC |
| *SULF1* | GAAGGAGAAGAGACGGCAGA | CAGAAAGATCCCAGGTTCCA |
| *SULF2* | CACTGGCAAGTACGTCCACAA | CTATTGAGGTACACCCCAAAGG |
| *HS6ST1* | GAGCTCAACGACCTGGACAT | TTGTACTGGTAGCGCTGCTG |
| *HS6ST2* | TACACTGGCGATGACTGGTC | GCGGTTGTTGGCTAGATTGT |
| *HS6ST3* | CTGGTGGGCTGCTATAACTTG | CACTCTGCAACAGGATGGTG |
| *HS3ST1* | AACGAGGTCCACTTCTTCGA | GCATCTGGCTGAGGTACCAG |
| *EED* | TTGCATTCGGCAATCAAGTT | GCAGCACCACATTTATGATGAG |
| *SUZ12* | CCGAGCACTGTGGTTGAGTA | AACTGCATCTGATGGTGGTG |
| *TRIM28* | TGTTTCCACCTGGACTGTCA | CCAGCAGTACACGCTCACAT |
| Negative (ChIP) | GCCACTTAGAAGTCGCAGGA | AGCAGACATACAACGGACGG |
| *SULF1* (ChIP) | TACGTGATATTGGCAGGGCA | CCCCTTAAGTAAATGATTTGCCG |

**Supplementary Table 3:**
HS disaccharide composition for A375 WT and *EZH2*$^{-/-}$ knockout cells

| Disaccharide Structure | | Abundance (% Total Disaccharide)[c] | |
|---|---|---|---|
| **Structure Code[a]** | **Unit Formula[b]** | **A375 WT (% Total HS)** | ***EZH2*$^{-/-}$ (% Total HS)** |
| D0A0 | ΔUA-GlcNAc | 53.87 ± 0.36 | 54.98 ± 0.67 |
| D0H6 | ΔUA-GlcNH$_2$6S | - | - |
| D2H0 | ΔUA2S-GlcNH$_2$ | 0.02 ± 0.01 | 0.03 ± 0.01 |
| D0S0 | ΔUA-GlcNS | 22.87 ± 0.46 | 22.88 ± 0.24 |
| D0A6 | ΔUA-GlcNAc6S | 4.97 ± 0.05 | 3.78 ± 0.04 |
| D2A0 | ΔUA2S-GlcNAc | 1.54 ± 0.03 | 1.56 ± 0.01 |
| D2H6 | ΔUA2S-GlcNH$_2$6S | 0.02 ± 0.02 | 0.17 ± 0.13 |
| D0S6 | ΔUA-GlcNS6S | 4.51 ± 0.27 | 4.19 ± 0.23 |
| D2S0 | ΔUA2S-GlcNS | 9.62 ± 0.12 | 9.44 ± 0.24 |
| D2A6 | ΔUA2S-GlcNAc6S | 0.02 ± 0.01 | 0.02 ± 0.01 |
| D2S6 | ΔUA2S-GlcNS6S | 2.55 ± 0.11 | 2.95 ± 0.16 |

[a] The disaccharide structure code is described in (Lawrence, et al. *Nat. Methods* 2008)
[b] ΔUA = 4,5-unsaturated uronic acid
[c] –, not detected

**Supplementary Table 4:**
HS sulfate groups per disaccharide of glucosamine units for A375 WT and *EZH2*$^{-/-}$ KO cells

| HS Sulfates/disaccharide | Abundance (% Total Disaccharide) | |
|---|---|---|
| | **A375 WT** | ***EZH2*$^{-/-}$** |
| 0 $SO_3$ | 53.87 ± 0.36 | 54.99 ± 0.66 |
| 1 $SO_3$ | 29.41 ± 0.38 | 28.24 ± 0.25 |
| 2 $SO_3$ | 14.17 ± 0.20 | 13.82 ± 0.37 |
| 3 $SO_3$ | 2.55 ± 0.11 | 2.95 ± 0.16 |

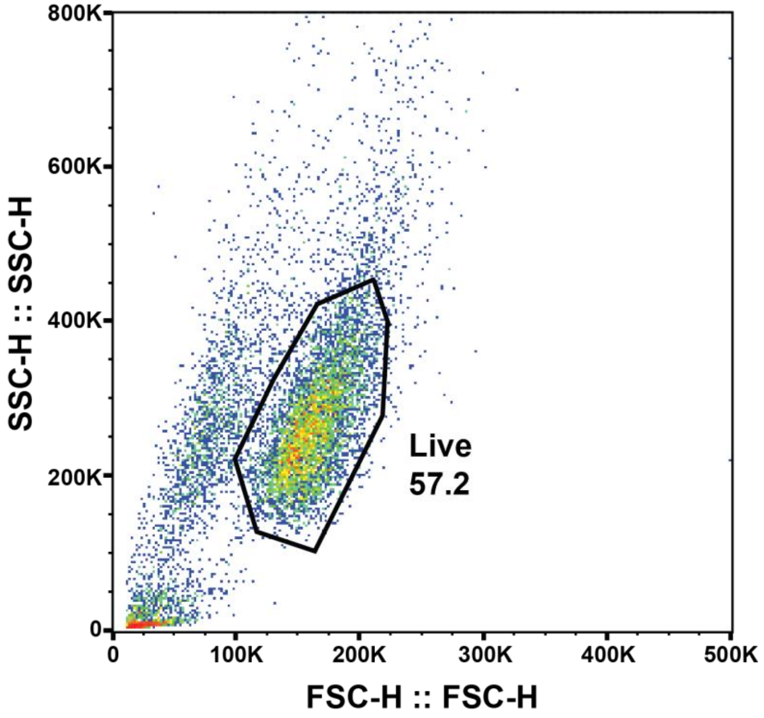


**Supplementary Figure 6. General flow cytometry gating strategy.** Cells were gated based on forward and side scattering for analysis of flow cytometry data.

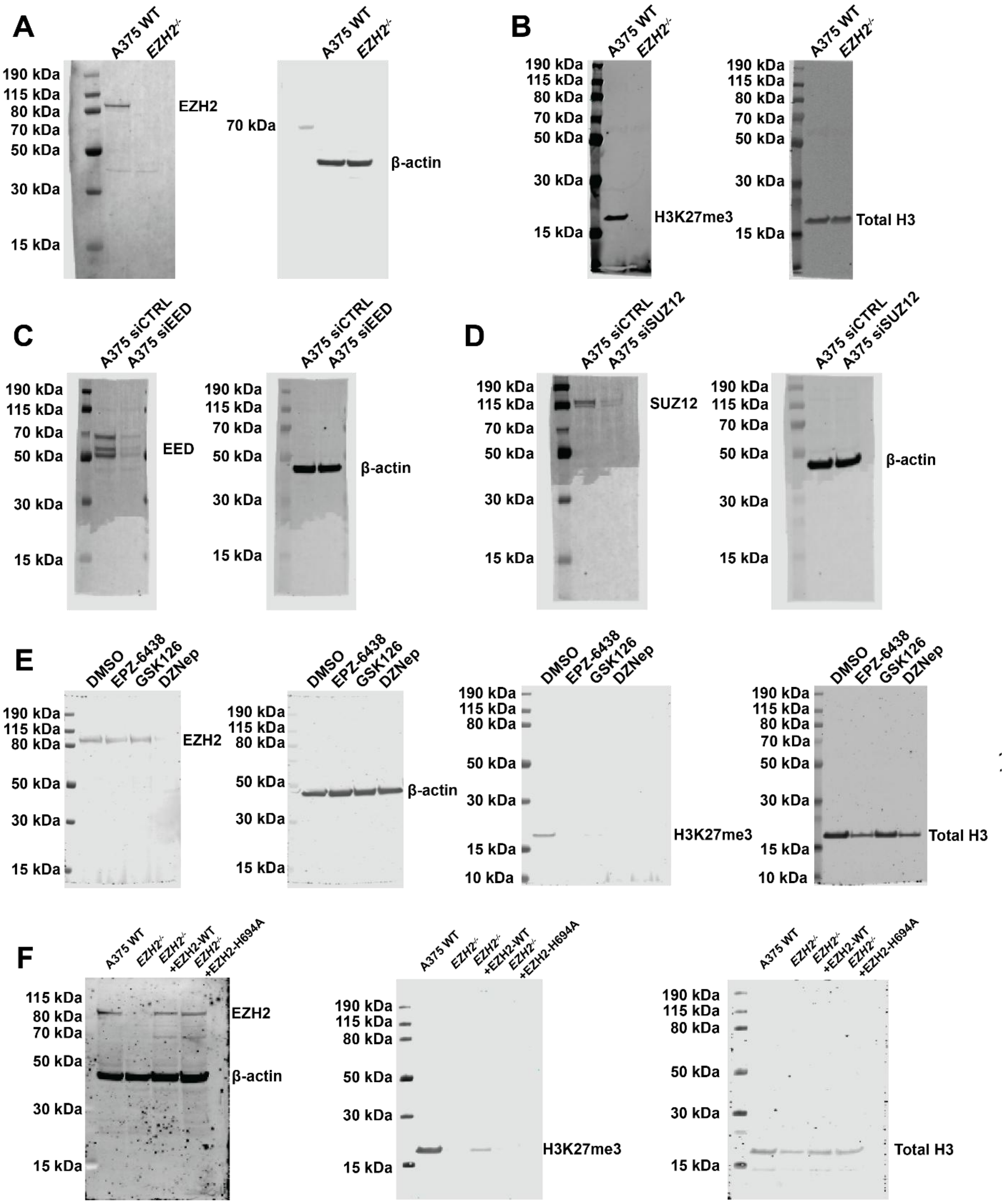


**Supplementary Figure 7. Source Data.** Uncropped western blot source images for **(A-B)** Figure 2A, **(C)** Figure 3A, **(D)** Figure 3C, **(E)** Figure 3F, and **(F)** Figure 3H.

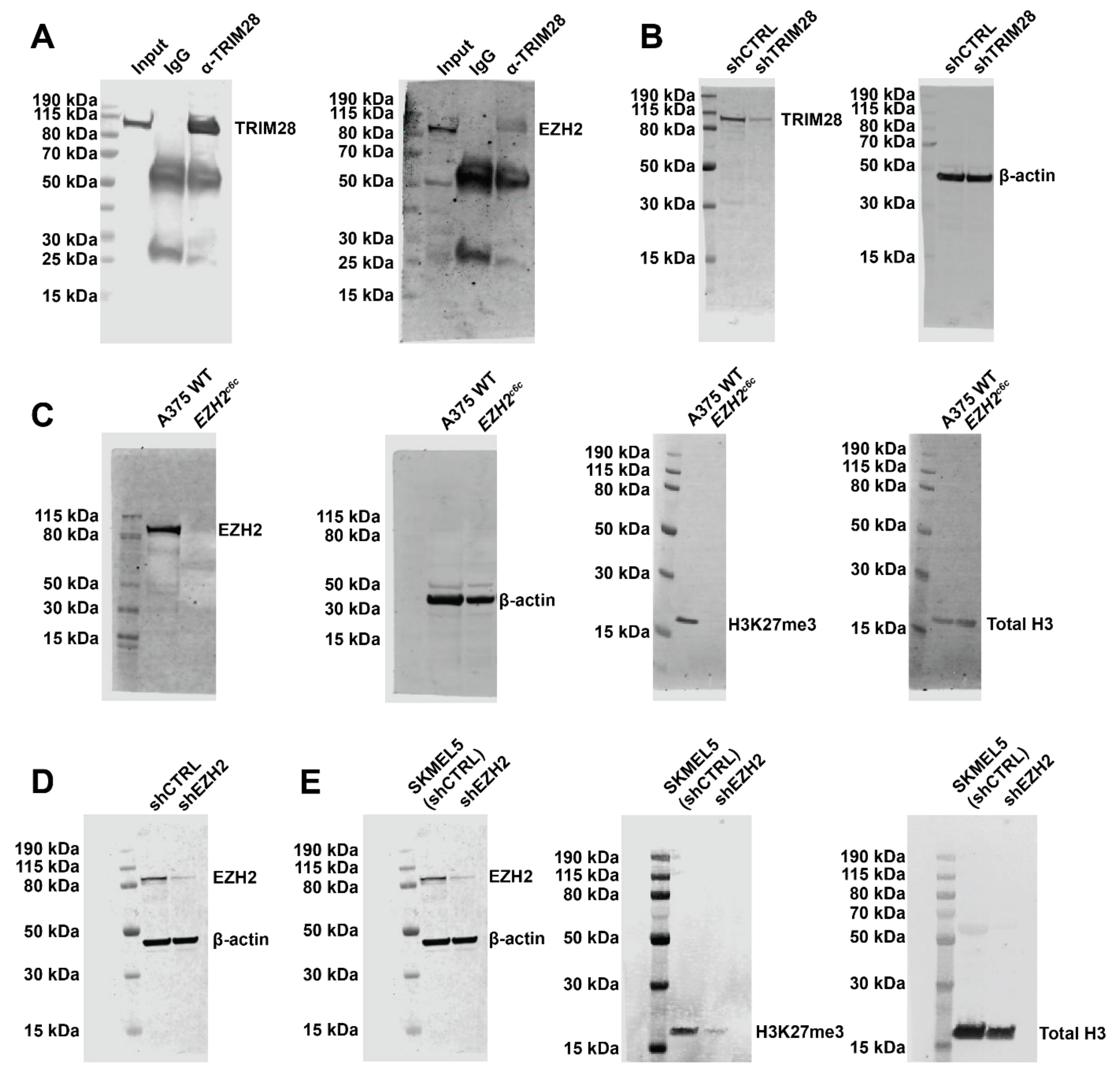


**Supplementary Figure 8. Source Data.** Uncropped western blot source images for **(A)** Figure 4B, **(B)** Figure 4C, **(C)** Suppl. Fig. 2E, **(D)** Suppl. Fig. 3A, and **(E)** Suppl. Fig. 4C.
